# Natural Antisense Transcript of *Period2, Per2AS*, regulates the amplitude of the mouse circadian clock

**DOI:** 10.1101/2020.07.27.222760

**Authors:** Rebecca A. Mosig, Allison N. Castaneda, Jacob C. Deslauriers, Landon P. Frazier, Kevin L. He, Naseem Maghzian, Aarati Pokharel, Camille T. Schrier, Lily Zhu, Nobuya Koike, John J. Tyson, Carla B. Green, Joseph S. Takahashi, Shihoko Kojima

## Abstract

Circadian transcriptome studies identified a novel transcript at the *Period2 (Per2)* locus, which we named *Per2AS. Per2AS* is a long non-coding RNA transcribed from the antisense strand of *Per2*, and is expressed rhythmically and anti-phasic to *Per2* mRNA. Previously, we mathematically tested the hypothesis that *Per2AS* and *Per2* mutually inhibit each other’s expression by forming a double negative feedback loop, and found that *Per2AS* expands the oscillatory domain. In this study, we have experimentally tested this prediction by perturbing the expression of *Per2AS* in mouse fibroblasts. We found that *Per2AS* represses *Per2* pre-transcriptionally *in cis* and regulates the amplitude of the circadian clock, but not period or phase. Unexpectedly, we also found that *Per2* positively regulates *Per2AS* post-transcriptionally, indicating that *Per2AS* and *Per2* form a single negative feedback loop. Because knock-down of *Per2* does not recapitulate the phenotypes of *Per2AS* perturbation and *Per2AS* also activates *Bmal1 in trans*, we propose that *Per2AS* regulates the amplitude of the circadian clock without producing a protein by rewiring the molecular clock circuit.

## Introduction

Eukaryotic genomes are pervasively transcribed, and non-protein coding portions of the genome dominate the transcriptional output of mammals (Bertone et al. 2004; Rosok and Sioud 2004; Carninci et al. 2005; Katayama et al. 2005; Sun et al. 2006). RNA species beyond mRNA are known as non-coding RNAs (ncRNAs) and have been categorized into many subtypes, such as ribosomal RNA (rRNA), transfer RNA (tRNA), small nucleolar RNAs (snoRNAs), small nuclear RNAs (snRNAs), microRNAs (miRNAs), circular RNAs (circRNAs), and long noncoding RNAs (lncRNAs) (Mercer et al. 2009; Panda et al. 2017). Despite their pervasive transcription, ncRNAs, particularly lncRNAs, were originally considered to be mere transcriptional noise and to lack defined functions. Poor evolutionary conservation in their primary sequences between species also raised concerns about their functional significance (Ponjavic et al. 2007; Guttman et al. 2009; Johnsson et al. 2014). More recent studies, however, have revealed an intriguing conservation of the genomic positions of lncRNAs (i.e., synteny) as well as conservation of their promoter and exon sequences, compared to intron or nontranscribed intergenic regions of protein-coding genes (Khaitovich et al. 2006; Pang et al. 2006; Yassour et al. 2010; Rhind et al. 2011; Derrien et al. 2012; Goodman et al. 2013; Anderson et al. 2016; Engreitz et al. 2016; Groff et al. 2016). These observations raise the possibility that some lncRNAs, if not all, are biologically relevant and have important and conserved functions. Indeed, a few dozen examples have highlighted the importance of lncRNAs in a variety of biological processes, such as X chromosome inactivation, imprinting, cell cycle regulation, and stem cell differentiation (Lee et al. 1999; Smilinich et al. 1999; Sleutels et al. 2002; Feng et al. 2006; Dinger et al. 2008; Morris et al. 2008; Zhao et al. 2008; Bond et al. 2009; Sopher et al. 2011; Modarresi et al. 2012).

Circadian rhythmicity, which modulates daily biochemical, physiological, and behavioral cycles, is a fundamental aspect of life on Earth. In mammals, essentially every cell is capable of generating circadian rhythms, and, within each cell, a set of clock genes forms a network of transcription-translation feedback loops that drive oscillations of approximately 24 hours (Lowrey and Takahashi 2004; Takahashi et al. 2008; Takahashi 2017). In one of these loops, the heterodimeric transcription activators (BMAL1/CLOCK and its paralogue BMAL1/NPAS2) activate transcription of the *Period (Per) 1-3* and *Cryptochrome (Cry) 1-2* genes, resulting in high levels of these transcripts. The resulting PER and CRY proteins then heterodimerize in the cytoplasm, translocate back to the nucleus, and interact with CLOCK/BMAL1 to inhibit transcription of *Per* and *Cry* genes. Subsequently, the PER/CRY repressor complex is degraded, and BMAL1/CLOCK can now activate a new cycle of transcription. In a second feedback loop, BMAL1/CLOCK activates the expression of orphan nuclear receptor genes *Rev-erbα/β* (*Nr1d1/2*). REV-ERB proteins, in turn, repress the expression of *Bmal1, Clock, Npas2, Cry1* and *Nfil3*. ROR proteins, also orphan nuclear receptors, recognize the same DNA motif as REV-ERB proteins, compete with their binding, and activate the expression of the same target genes. In the last loop, BMAL1/CLOCK activates the expression of *Dbp*, while REV/ROR proteins activate and inhibit the expression of *Nfil3*, respectively. Both DBP and NFIL3 are transcription factors: DBP activates while NFIL3 represses the transcription of target genes, such as *Rev-erbs, Rors*, and *Pers*. These feedback loops constitute the molecular mechanism, as currently understood, of circadian rhythms in mammals (Takahashi 2017).

Several circadian transcriptome studies discovered a new RNA molecule, which we named *Per2AS*, that is transcribed within the *Per2* locus in mouse liver, lung, kidney and adrenal gland (Koike et al. 2012; Menet et al. 2012; Vollmers et al. 2012; Fang et al. 2014; Zhang et al. 2014). In mammals, 25-40% of protein-coding genes have antisense transcript partners (Rosok and Sioud 2004; Katayama et al. 2005; Sun et al. 2006), and some antisense transcripts have been shown to exert functions in a variety of processes, such as cell cycle regulation, genome imprinting, immune response, neuronal function and cardiac function (Faghihi and Wahlestedt 2009; Khorkova et al. 2014; Wanowska et al. 2018). However, the physiological roles of most antisense transcripts remain uncertain.

In an earlier publication, we constructed mathematical models of *Per2-Per2AS* interactions, assuming that *Per2AS* and *Per2* mutually inhibit each other’s expression in either pre-transcriptional (i.e., transcriptional interference) or post-transcriptional (i.e., *in trans* and degradation of RNA duplexes), or a combination of effects. The model predicted that all three mechanisms are consistent with the basic molecular details of circadian rhythms in mouse, but the pre-transcriptional model gives a more robust account of the circadian, anti-phasic oscillations of *Per2* and *Per2AS*, compared to the post-transcriptional model (Battogtokh et al. 2018). To test predictions of these models, we have experimentally perturbed the expression of *Per2AS* both pre- and post-transcriptionally in this study. We found that *Per2AS* pre-transcriptionally represses *Per2*, however, *Per2* positively regulates *Per2AS* post-transcriptionally. We also found that *Per2AS* regulates the amplitude of the circadian clock, although this effect is not solely dependent on its interaction with *Per2*. Overall, we propose that *Per2AS* serves as an important regulatory molecule not only to regulate the amplitude, but also to limit the level of *Per2* within the oscillatory range. These conclusions are significant because *Per2* is the only core clock gene for which the abundance, rhythmicity, and the phase of its expression are critical to maintain circadian rhythmicity in mouse (Chen et al. 2009).

## Results

### Characterization of *Per2AS*

Recent circadian transcriptome studies have identified a novel transcript, *Per2AS*, at the *Per2* locus that is transcribed from the opposite strand, and expressed anti-phasic to the sense *Per2* transcript in mouse liver (Koike et al. 2012; Menet et al. 2012; Vollmers et al. 2012; Fang et al. 2014; Zhang et al. 2014). Our quantitative PCR (qPCR) analyses demonstrated that the *Per2AS* is indeed expressed rhythmically and anti-phasic to *Per2* mRNA in mouse liver, and it peaks at ZT 4 (Zeitgeber time, where ZT 0 and ZT 12 are defined as time of lights on and lights off, respectively) (Fig. 1A). Rhythmic and anti-phasic expression patterns of *Per2AS* and *Per2* have been observed, as well, in NIH3T3 cells (*Bmal1*-luc) and mouse embryonic fibroblasts (MEFs) derived from *PER2::LUCIFERASE* knock-in mouse (Fig. 1A) (Yoo et al. 2004).

**Fig. 1.**
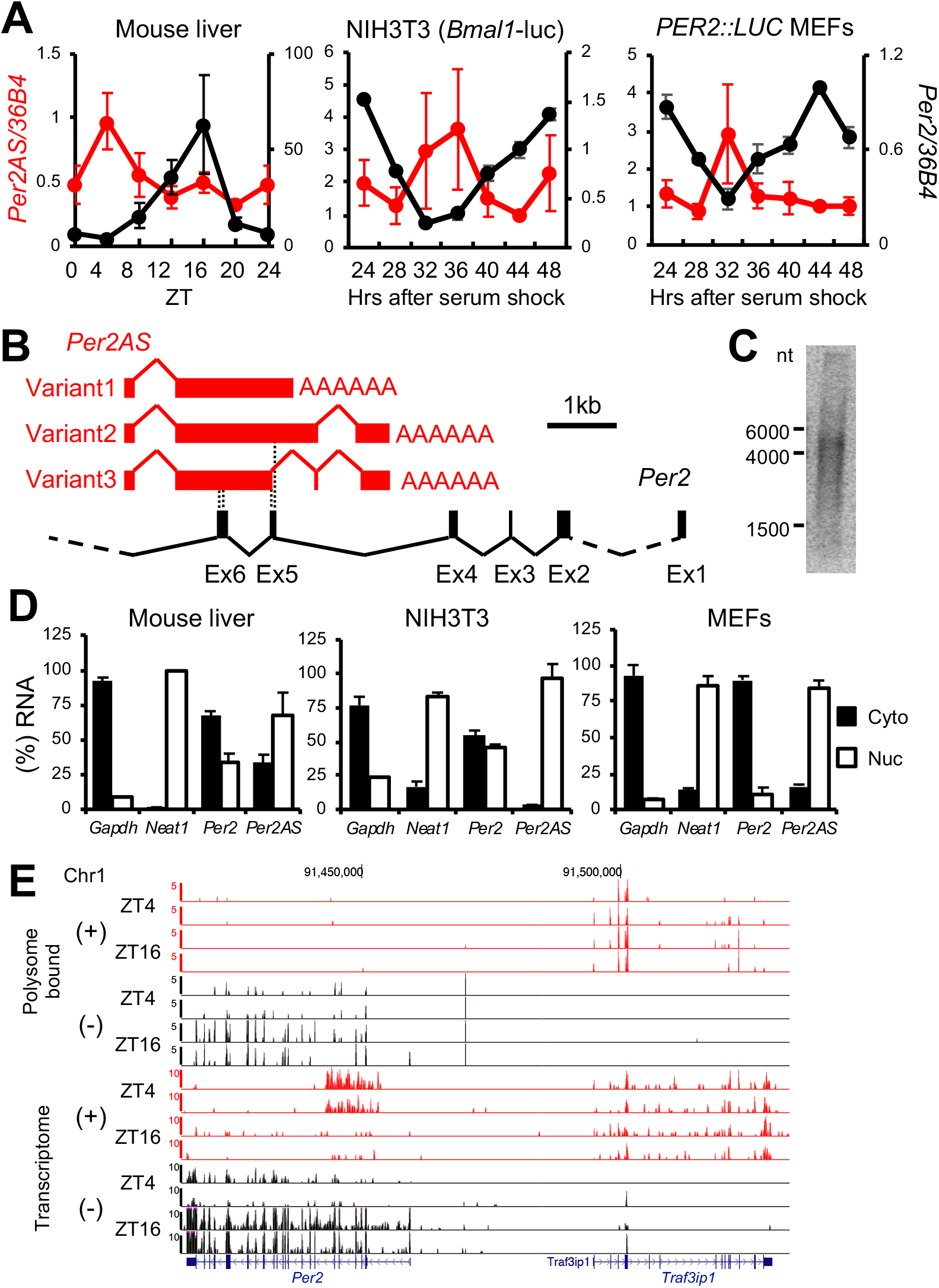
Characterization of *Per2AS*. A) Relative RNA expression of *Per2AS* (red) and *Per2* (black) in mouse liver, *Bmal1-luc* (NIH3T3), and *PER2::LUC* MEFs. The expression levels of *Per2AS* at 44 hrs after serum shock were set to 1 in *Bmal1-luc* and *PER2::LUC* MEFs. B) Genomic structure of *Per2AS* (red) in relation to *Per2* (black). Dotted lines indicate the regions where the exons of *Per2* and *Per2AS* overlap. C) Northern Blot analysis of *Per2AS*. PolyA^+^ enriched RNA was extracted from mouse liver at ZT 4 and probed with *Per2AS* sequence (1-1812nt that are shared in all the variants, depicted in B. Numbers on the left represent the size of RNA markers. D) Subcellular localization of *Per2AS* and *Per2* RNAs in mouse liver at ZT 4 (left), NIH3T3 cells (middle) and MEFs (right). *Neat1;* nuclear RNA, *Gapdh;* cytoplasmic RNA serving as controls. E) Genome browser view on chromosome 1 for both ribosome-bound transcripts (top) and ribosome-depleted RNAs (bottom). Data analyzed are from (Janich et al. 2015). All the data represent Mean ± SEM (n=2-5).

*Per2AS* is spliced and polyadenylated, similar to mRNAs (Figs. 1B, S1A). Rapid Amplification of cDNA Ends (RACE) analysis revealed that the transcription start site (TSS) of *Per2AS* is located in intron 6 of *Per2* (Fig. 1B), which is consistent with the most 5’-signals of *Per2AS* (Fig. S1A) (Koike et al. 2012; Menet et al. 2012; Zhang et al. 2014). Strong and rhythmic recruitment of RNAPII-Ser5P (RNA polymerase II whose serine 5 in the C-terminal domain is phosphorylated) has been observed just upstream of *Per2AS* TSS, indicating an active and rhythmic initiation of transcription at this site (Phatnani and Greenleaf 2006) (Fig. S1A). RNAPII’s recruitment pattern also coincides with the expression pattern of *Per2AS* (Fig. S1B), indicating that *Per2AS* is transcribed by RNAPII.

*Per2AS* consists of at least three variants ranging in size from ~2000 to 3500 nt (Fig. 1B), while Northern Blot analysis yielded a strong signal around ~5000 nt (Fig. 1C). Given that the signal from Northern Blot was smeared, ranging from ~1500 to 5000 nt (Fig. 1C), and the *Per2AS* signals from circadian transcriptome studies were detected in a 10^+^ kb range (Fig. S1A) (Koike et al. 2012; Menet et al. 2012; Zhang et al. 2014), *Per2AS* presumably has many variants that are different in splicing patterns and transcription termination sites.

All three *Per2AS* variants that we detected by RACE have a potential to encode a small protein (the longest ORF common to all three is 291 nt; 97 amino acids); however, the potential polypeptide has no sequence similarity to any existing or predicted polypeptides in the GenBank protein database or the Conserved Domain Database. Furthermore, the Coding Potential Calculator (http://cpc.cbi.pku.edu.cn/) predicts that *Per2AS* does not encode a protein, as the coding score of *Per2AS* is much smaller (from −0.97 to −0.99 depending on variant) than that of *Per2* (17.53). Furthermore, *Per2AS* transcripts are predominantly detected in the nucleus in mouse liver, NIH3T3 cells, and MEFs (Fig. 1D), similar to *Neat1*, a lncRNA known to localize in the nucleus (Clemson et al. 2009). In contrast, protein-coding transcripts, such as *Gapdh* and *Per2*, were localized largely in the cytoplasm in order for them to be translated (Fig. 1D). In addition, *Per2AS* is not bound to polysomes and therefore not actively translated, in contrast to *Per2* that shows strong signals both in ribosome profiling and transcriptome data and those signals are higher at ZT16 compared to ZT4 (Fig. 1E, Fig. S1C) (Atger et al. 2015; Janich et al. 2015). The neighboring protein-coding transcript, *Traf3ip1*, that is transcribed from the same strand as *Per2AS*, also showed a clear signal both in ribosome profiling and transcriptome data, eliminating the possibility that lack of *Per2AS* signals in the polysome-bound fraction is due to its strand. These data collectively indicate that *Per2AS* is a long non-coding antisense transcript and does not produce a protein.

### Per2AS regulates Per2 and the amplitude of the circadian clock

To understand the functional relevance of *Per2AS* in the mammalian circadian clock, we first used CRISPR technology to perturb *Per2AS* expression. We targeted the putative *Per2AS* promoter region, defined by the strong and rhythmic recruitment of RNAPII-Ser5P spanning approximately 900 bp (Fig. S1A-B). We introduced CRISPR mutagenesis in two independent cell lines: MEFs from *PER2::LUCIFERASE (PER2::LUC)* knock-in reporter mice, in which a luciferase gene is fused to the 3’-end of the endogenous *Per2* gene (Yoo et al. 2004), and *Bmal1-* luc, an NIH3T3-derived luciferase-reporter cell line that has been stably transfected by a luciferase gene driven by the *Bmal1* promoter (Morf et al. 2012). Following single cell sorting, we successfully isolated two *PER2::LUC MEF* (5D8 and 6F8) and one *Bmal1-luc* (mut8) mutant clones that had similar but distinct mutations at approximately 260 bp upstream of *Per2AS* TSS (Fig. S1D). Because these cell lines were near tetraploid on average (Fig. S1E), we not only identified the types of mutations, but also calculated the frequency of each mutant allele in each mutant (Fig. S1D). All the alleles were mutated in *PER2::LUC* 5D8 and 6F8, with which various deletions were detected (i.e., 11bp, 3bp, 2bp, and 1bp). Meanwhile in *Bmal1-luc* (mut8), only 29% of alleles had a mutation (6 bp deletion) and the remaining 71% were intact (Fig. S1D). We also targeted five other regions within the *Per2AS* promoter, including TSS and the TATA-box-like sequence (TATAATCAA) located 63 bp upstream of the TSS; however, we were unable to isolate mutant clones, probably because these targeted regions were essential for cell survival or could not be easily accessed by the CRISPR machinery.

In these mutant cell lines, the level of *Per2AS* was upregulated to 138% (5D8) and 325% (6F8) compared to the control (parental) cell line, as opposed to our expectations that *Per2AS* would be downregulated. Nonetheless, the level of *Per2* was reduced to 57% (5D8) and 70% (6F8), respectively (Fig. 2B). Similarly, in *Bmal1-luc* mut8, the *Per2AS* level was upregulated to 145%, while the *Per2* level was reduced to 76% (Fig. 2F). These data indicate that the DNA region approximately 260bp upstream of the *Per2AS* TSS is important for *Per2AS* expression and that *Per2AS* represses *Per2*. Some long non-coding antisense transcripts modulate the expression of not only their target sense gene, but also neighboring genes (Halley et al. 2014; Villegas et al. 2014). However, this was not the case for *Per2AS*, as the levels of *Ilkap, Hes6* and *Traf3ip1*, the three closest genes, remained unchanged in all of our mutants (Fig. S1F-G).

**Fig. 2.**
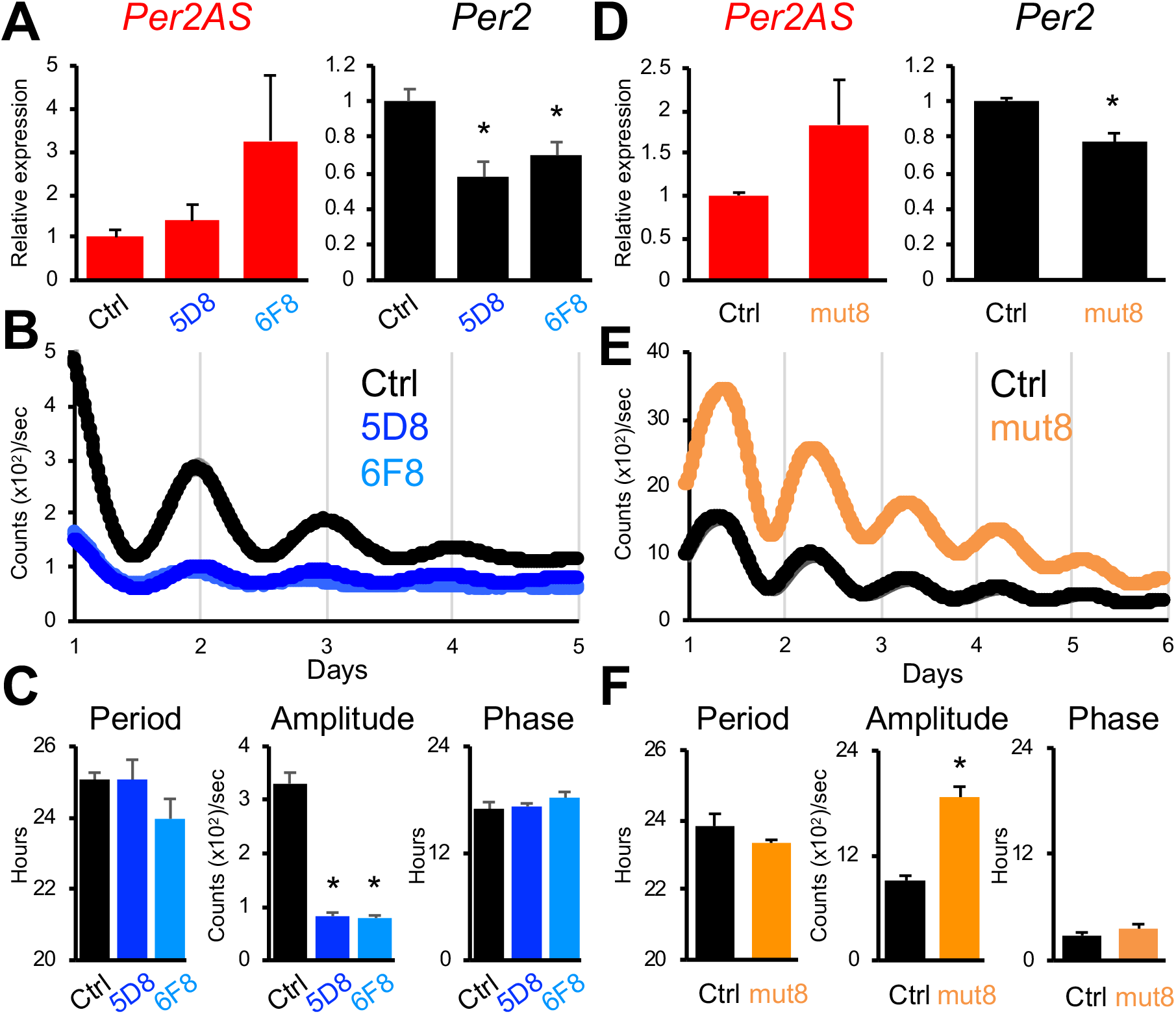
*Per2AS* mutants alter *Per2* and the amplitude of the circadian clock. A, D) Relative expression levels of *Per2AS* (red), and *Per2* (black) in the *Per2AS* mutant cell lines of *PER2::LUC* MEFs (n=4-6) (A) or *Bmal1-luc* (D) (n=13). B, E) Bioluminescent output of the mutant cell lines of *PER2::LUC* MEFs (WT: n=12, 5D8: n=4, 6F8: n=6) (B) or *Bmal1-luc* (WT: n=9, mut8: n=9) (E). Bold lines represent the mean, while shaded areas represent the SEM. *C, F*) Period (left), amplitude (middle), and phase (right) of bioluminescent output calculated from Fig. 2B (*PER2::LUC* MEFs) or 2E *(Bmal1-luc)*. All the data represent Mean ± SEM. *; p<0.05 (Student t-test).

When we monitored the bioluminescent output from these mutant cells, the luminescent levels of both 5D8 and 6F8 were lower and their rhythmicity was less robust compared to the control (Fig. 2B, S2A-B). Quantification analyses further revealed that the amplitude of 5D8 and 6F8 was decreased to 25% of the control, whereas the period and phase remained unchanged (Fig. 2C). In contrast, the bioluminescence signal from *Bmal1-luc* mut8 was markedly higher (Fig. 2E, S2C-D), and the amplitude was increased to 204%, while the period and phase remained unchanged, compared to the control cell line (Fig. 2F). These data indicate that *Per2AS* regulates *Per2*, as well as the amplitude of the circadian clock. It is unlikely the observed phenotypes are due to an off-target effect, because mutants from two independent cell lines (three independent clones) with similar but distinct mutations all led to the same phenotype.

To further characterize the *Per2AS* mutants as well as to gain insights into the underlying mechanisms of circadian amplitude regulation, we next measured the mRNA expression patterns of 13 core clock genes after synchronizing these cells by serum shock (Balsalobre et al. 1998). We found that the mRNA expression of *Bmal1, Cry1*, and *Rorα* were elevated, while the expression of *Per2* was decreased in all three mutants (Fig. 3). The up-regulation of *Bmal1* mRNA level was consistent with the increased bioluminescence levels in *Bmal1-luc* cells (Fig. 2E). The mRNA expression of *Nfil3* was up-regulated in *PER2::LUC* 5D8 and *Bmal1-luc* mut8, while that of *Cry2* and *Dbp* was down-regulated only in *PER2::LUC* mutants (5D8 and 6F8) (Fig. 3). Interestingly, the expression of *Npas2* was down-regulated in the *PER2::LUC* mutants but up-regulated in the *Bmal1-luc* mutant. No significant changes were observed for *Rev-erbα* and *Rev-erbβ* (Fig. 3). These data suggest that *Bmal1, Cry1, Rora, Per2* (changed in all three mutants), but not *Rev-erbα/β*, are involved in the *Per2AS*-mediated amplitude regulation.

**Fig. 3.**
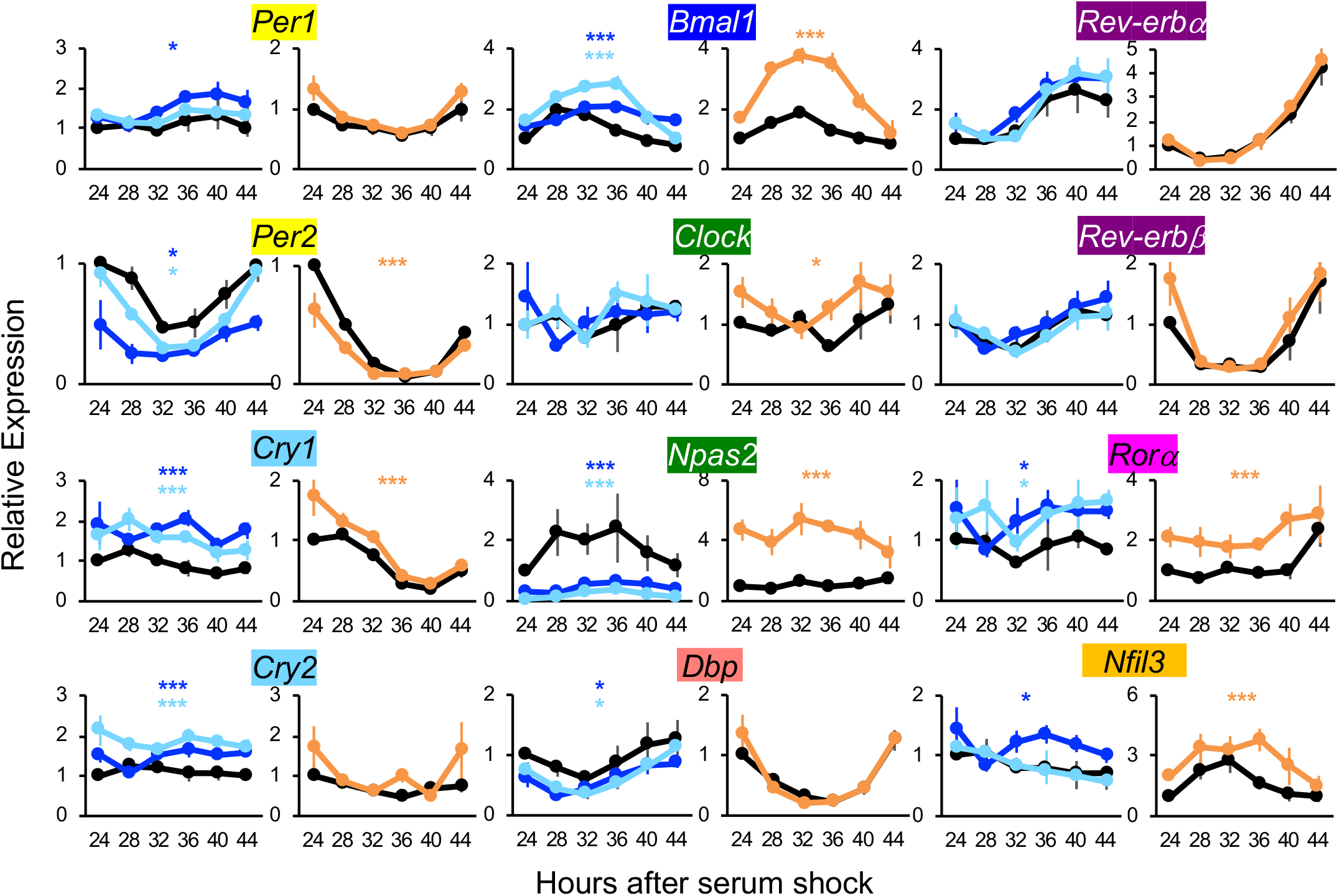
Relative expression levels of core clock genes in the *Per2AS* mutant cell lines. All the data represent Mean ± SEM. *PER2::LUC* MEFs (WT: n=3, 5D8: n=3, 6F8: n=3) (B) or *Bmal1-luc* (WT: n=3, mut8: n=3) (E). *; p<0.05, ***; p<0.005 (Two-way ANOVA).

### Transcripts of Per2AS do not play a major role in the circadian clock machinery

In contrast to coding genes whose functional unit is an encoded protein, the functional unit of lncRNAs is often unknown. It can be the transcript itself (i.e., RNA molecule and/or the small ORF embedded in the sequence that has the potential to encode a protein) that regulates target mRNAs post-transcriptionally by affecting splicing, mRNA stability, mRNA localization, or epigenetic marks (Wight and Werner 2013; Khorkova et al. 2014). It can also be the very act of transcription, in which transcription of one strand suppresses transcription of another *in cis*, a process called transcription interference (Wight and Werner 2013; Khorkova et al. 2014).

To distinguish whether the functional unit of *Per2AS* is either a transcript (i.e., post-transcriptional model) or the act of transcription (i.e., pre-transcriptional model) (Battogtokh et al. 2018), we post-transcriptionally decreased the level of *Per2AS* using “gapmers”, chimeric antisense nucleotides containing modified nucleic acid residues to induce RNase H-mediated degradation of nuclear-retained RNA (Lee et al. 2012), as the majority of *Per2AS* RNA remains in the nucleus in fibroblasts (Fig. 1D) and RISC-mediated RNA cleavage triggered by siRNAs occurs mainly in the cytoplasm (Carthew and Sontheimer 2009). We also targeted gapmers to exon 1 of *Per2AS* (i.e., intron 6 of *Per2*) that was shared by all three variants (Fig. 1B). Gapmers 5 and 8 successfully reduced the level of *Per2AS* to 61% in *Bmal1-luc* cells (Fig. 4A); however, the level of *Per2* remained unchanged (Fig. 4A). Even though gapmers can not only induce RNA duplex-mediated degradation but also premature transcriptional termination of target mRNAs leading to reduced transcriptional activity and thereby confounding the interpretation of the results (Lee and Mendell 2020), this is unlikely the case for *Per2AS* gapmers, as the level of *Per2* is unaffected. Unexpectedly, however, gapmer-mediated *Per2AS* knock-down led to an approximately 20% reduction in the *Bmal1* level in *Bmal1-luc* cells (Fig. 4A).

**Fig. 4.**
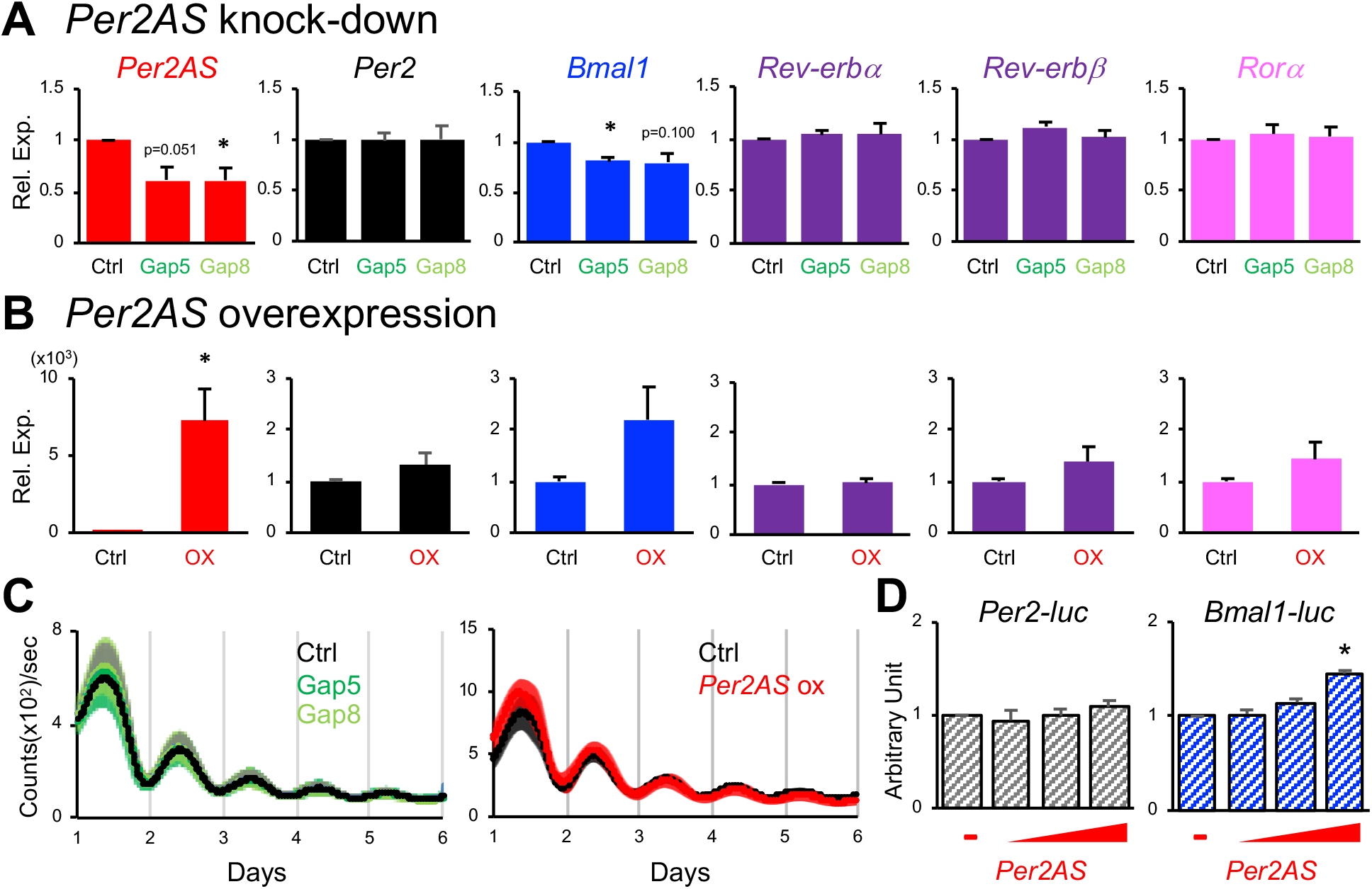
*Per2AS* transcripts regulate *Bmal1* but not *Per2* in *Bmal1-luc* cells. A) Relative expression levels of core clock genes normalized by *36B4* upon *Per2AS* knock-down (n=5). B) Relative expression levels of core clock genes normalized by *36B4* upon *Per2AS* overexpression (n=7-8). All the data represent Mean ± SEM. *; p<0.05. We excluded the samples that had less than 5000 fold induction of *Per2AS*. C) Luciferase activity from *Per2-luc* or *Bmal1-luc* cotransfected with an increasing amount of *Per2AS* (variant2)-expressing plasmid. D) Bioluminescent output from *Bmal1-luc* cells upon knock-down (n=8) or overexpression (n=4) of *Per2AS*. Bold lines represent the mean, while shaded areas represent the SEM.

We also post-transcriptionally overexpressed *Per2AS* using variant 2 (i.e., the longest variant) (Fig. 1B) in *Bmal1-luc* cells. This led to a marked increase in the *Per2AS* level by ~75,000-fold. Nevertheless, the *Per2* level remained unchanged, while the *Bmal1* level showed an approximate two-fold increase (Fig. 4B). Knock-down or overexpression of *Per2AS* did not alter period, phase, or amplitude of bioluminescence output in *Bmal1-luc* cells (Fig. 4C). The change in *Bmal1* level was not due to comparable changes in the levels of *Rev-erbα/β* or *Rorα*, transcription repressor and activators of *Bmal1*, respectively (Sato et al. 2004) (Fig. 4A-B). Rather, *Per2AS* appears to directly regulate the *Bmal1* transcription, as the increasing amount of *Per2AS* RNAs led to upregulation of reporter activity of *Bmal1-luc*, but not *Per2-luc* (Fig. 4D). Interestingly, however, the changes of *Bmal1* by *Per2AS* RNA was also cell type-specific, as it was not observed in *PER2::LUC* MEFs, even though the efficiency of *Per2AS* knock-down was higher (gapmer 5: 63%, gapmer 8: 60%) in *PER2::LUC* MEFs, compared to *Bmal1-luc* cells (both gapmer 5 and 8: 39%) (Fig. S3). Overall, these data indicate that the transcript of *Per2AS* has little effect on the circadian control system (i.e., that its effects are not post-transcriptional). However, we also observed a small effect of *Per2AS* on *Bmal1* at least in *Bmal1-luc* cells, suggesting that *Per2AS* post-transcriptionally regulates *Bmal1* in a cell-specific manner.

### Per2 knock-down downregulates Per2AS and does not replicate the phenotypes of Per2AS CRISPRI mutants

If *Per2AS* and *Per2* indeed form a double negative feedback loop and mutually inhibit each other’s expression, then *Per2* mRNA knock-down would result in up-regulation of *Per2AS* level in the post-transcriptional model but no changes in the pre-transcriptional model. To test this, we first reduced the level of *Per2* post-transcriptionally using shRNAs. Our *Per2* knockdown successfully reduced the level of *Per2* to 65% in AML12 cells, a mouse hepatocyte cell line, and 45% in *Bmal1-luc* cells (Fig. 5A, F). *Per2* knock-down also led to a decrease in the amplitude of bioluminescence output in *PER2::LUC* MEFs but not in *Bmal1-luc* cells (Fig. 5B-D). Period remained unchanged upon *Per2* knock-down in both cells, despite that previous studies in murine fibroblast and hepatocyte cell lines reported that circadian period became shorter upon *Per2* knock-down (Ramanathan et al. 2014). The residual *Per2* level is higher in our system compared to those reports (45% vs 20%), and this could have contributed to the difference observed in our study. Nevertheless, *Per2* knock-down resulted in a decrease of the *Per2AS* level to 55% (Fig. 5A), contrary to either of our expectations (i.e., pre-transcriptional or post-transcriptional interference).

**Fig. 5.**
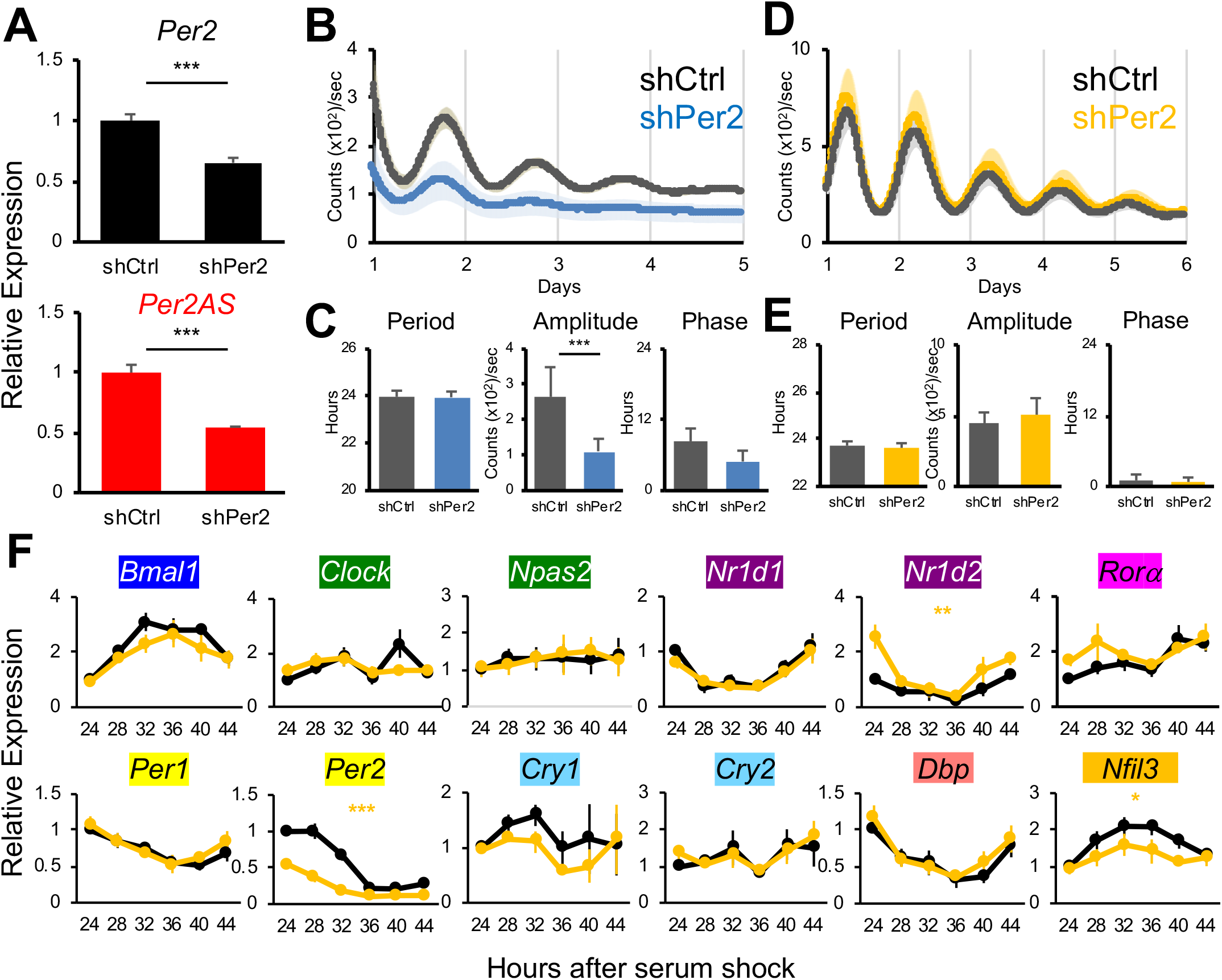
*Per2* knock-down does not replicate the phenotypes of *Per2AS* mutants. (A, B) Relative expression levels of *Per2* (A) and *Per2AS* (B) in the AML12 mouse hepatocytes (n=4-5). (B, D) Bioluminescence output from *PER2::LUC* MEFs (B) (n=12) or *Bmal1-luc* (D) (n=14) upon *Per2* knock-down. (C, E) Period (left), amplitude (middle), and phase (right) of bioluminescent output calculated from Fig. 5B (*PER2::LUC* MEFs) or 5D *(Bmal1-luc)*. ***; p<0.005 (Students’ t-test). (F) Relative expression levels of core clock genes in *Bmal1-luc* cells upon *Per2* knock-down (n=3-5). All the data represent Mean ± SEM. *; p<0.05, **; p<0.01, ***; p<0.005 (Two-way ANOVA).

We also measured the expression patterns of the core clock genes upon *Per2* knock-down in *Bmal1-luc* cells to evaluate whether the changes observed in the *Per2AS* mutant cells (Fig. 3) were mediated entirely through *Per2AS*’s effect on *Per2. Per2* knock-down did not result in changes of *Bmal1* mRNA levels (Fig. 5F), consistent with the bioluminescent output from *Bmal1-luc* cells (Fig. 5D), but inconsistent with our observations in *Bmal1-luc* mut8 cells, in which *Bmal1-luc* bioluminescence and *Bmal1* mRNA level were both significantly increased (Fig. 2E, 3). In addition, changes in mRNA expression patterns were observed only for *Nr1d2* and *Nfil3* upon *Per2* knock-down (Fig. 5F), in contrast to the changes in the mRNA expression patterns of *Cry1, Bmal1, Clock, Npas2, Rora*, and *Nfil3* observed in *Bmal1-luc* mut8 cells (Fig. 3). Interestingly, we did not observe any changes in mRNA levels of the core clock genes upon *Per2* knock-down in AML12 cells (Fig. S4A), in which the level of *Per2AS* is approximately 100-fold higher compared to *Bmal1-luc* cells (Fig. S4B). As the level of *Per2* was lower in *Bmal1-luc* with *Per2* knock-down (45%) compared to *Bmal1-luc* mut8 cells (75%), it is highly unlikely that changes in the mRNA expression patterns observed in *Bmal1-luc* mut8 cells (Fig. 3) are solely due to the *Per2AS*’s effect on *Per2*. Rather, these data support the idea that *Per2AS* has a distinct role in the mammalian circadian system, independent of *Per2*.

## Discussion

There is only limited experimental evidence about the functions and regulatory mechanisms of antisense transcripts, particularly in mammals. Our study contributes to an understanding of the possible functional roles of antisense transcripts by demonstrating that *Per2AS*, a natural antisense transcript to a core clock gene *Per2*, regulates the amplitude of the circadian clock. Although the precise regulatory mechanism remains unclear, we think *Per2AS* does not solely rely on *Per2* for its functions. *Per2* knock-down leads to decreased levels of RORE-controlled genes, such as *Bmal1* and *Nfil3* (Shearman et al. 2000; Schmutz et al. 2010; Ramanathan et al. 2014); however, *Per2AS* mutants that upregulate antisense transcription do not exhibit these changes. Of potential interest in understanding the amplitude-regulatory mechanism of *Per2AS* would be *Rora*, as *Rora* is one of the core clock genes whose expression is elevated in all the *Per2AS* mutants in our study (Fig. 3). Interestingly, *Ror* genes have been shown to increase the circadian amplitude of molecular and behavioral rhythms (He et al. 2016). Furthermore, our recent analysis suggested that *Rorc* and *Per2AS* are potential amplitude regulators of circadian transcriptome output, as the levels of *Rorc* and *Per2AS* correlate with the percentage of rhythmic transcripts in various mouse tissues (Littleton and Kojima 2020). Notably, existing *PER2* null animals all lack the *Per2AS* locus (Zheng et al. 1999; Bae et al. 2001), and thus complicate the interpretation of the data from these animals in understanding whether the observed phenotypes are solely due to *Per2*, or a combined effect of *Per2AS* and *Per2*.

The experimental observations from this study bear on the assumptions underlying our mathematical models of *Per2-Per2AS* interactions (Battogtokh et al. 2018). In building our models, we considered that *Per2AS* and *Per2* mutually inhibit each other’s abundance either pre-transcriptionally or post-transcriptionally, or a combination of both effects (Battogtokh et al. 2018). Our experimental interrogation clearly demonstrated that *Per2AS* represses *Per2* (Fig. 2A, D), presumably via a pre-transcriptional mechanism, as the knock-down or overexpression of *Per2AS* transcripts did not alter the level of *Per2* mRNAs (Fig. 4, 6). Contrary to our expectation, however, *Per2* positively regulates *Per2AS*, as the knock-down of *Per2* led to downregulation (rather than upregulation) of *Per2AS* (Fig. 5A). This effect is post-transcriptional, because the level of *Per2* pre-mRNA remained unchanged upon *Per2* knock-down (Fig. S4C). We think it is unlikely that *Per2* RNA and *Per2AS* RNA post-transcriptionally form an RNA duplex and stabilize each other, because (1) the expression of *Per2AS* and *Per2* are anti-phasic in at least some tissues (Fig. 1A) (Zhang et al. 2014), (2) *Per2* is ~25x more abundant than *Per2AS* (Fig. 1A) (Koike et al. 2012), (3) *Per2* predominantly localizes in the cytoplasm while *Per2AS* localizes in the nucleus (Fig. 1D), and (4) *Per2AS* consists of many variants and their nucleotide sequences vary (Fig. 2E). Rather we favor the hypothesis that PER2 indirectly or directly regulates *Per2AS* transcription. In fact, REV-ERBα/β and BMAL1/CLOCK/PER1/PER2/CRY2 are recruited to the vicinity of the *Per2AS* TSS (Koike et al. 2012), potentially involved in regulating *Per2AS* transcription.

**Fig. 6.**
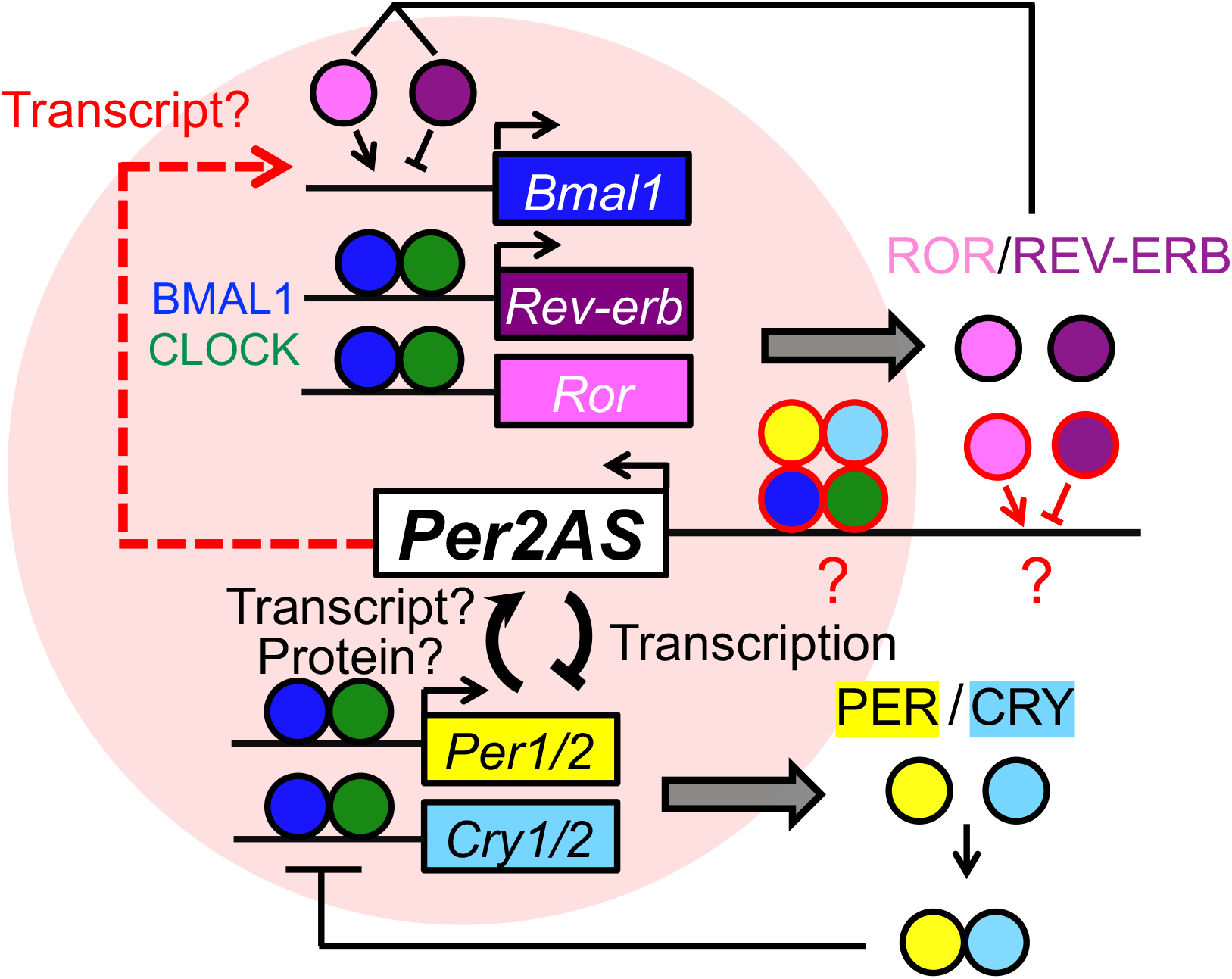
Putative model for the circadian molecular circuit including *Per2AS*. *Per2AS* and *Per2* form a double negative feedback and inhibit each other’s expression presumably pre-transcriptionally. *Per2AS* can also activate the *Bmal1* expression post-transcriptionally (i.e., *in trans*) in a tissue or cell-specific manner.

These experimental observations clearly indicate that our mathematical models need to be revised to better understand the functions and regulatory mechanisms of *Per2AS*. Specifically, we need to alter the description of the *Per2AS-Per2* relationship, include the effect of *Per2AS* on *Bmal1*, and add the potential effect of circadian transcription factors on *Per2AS* transcription (Fig. 6). Nevertheless, our original model predicted that the region of circadian oscillations is greatest in the pre-transcriptional model (Battogtokh et al. 2018). This conclusion is also true in a simpler mathematical model, involving only the core negative feedback loop, as we show in Fig. S5.

Our mathematical models were also constrained by the requirement that the expression of *Per2AS* and *Per2* are both rhythmic and anti-phasic (180° out of phase). These patterns have been observed not only in liver but also in adrenal gland, lung and kidney (Zhang et al. 2014). Analysis of mouse ENCODE datasets demonstrated that *Per2AS* is also expressed in many other tissues, such as genital and subcutaneous fat pad, bladder, kidney, colon, duodenum, large and small intestines, stomach, lung, and potentially in mammary gland, ovary and testis (Consortium 2012; Davis et al. 2018) (Fig. S6A), although it is unclear whether *Per2AS* expression is rhythmic and antiphasic to *Per2* in these tissues. A *Per2AS* signal has not been detected in the central nervous system and a few other peripheral tissues including heart, placenta, spleen, and thymus (Consortium 2012; Davis et al. 2018) (Fig. S6A). These results could be due to tissue sampling times, i.e., little or no *Per2AS* may have been expressed when these tissues were harvested, or the *Per2AS* level was under detection threshold, as the expression of lncRNAs is generally low (more than 10-fold lower than sense transcripts on average) and transcriptome analyses sometimes lack the sensitivity to detect all lncRNAs (He et al. 2008; Faghihi and Wahlestedt 2009; Xu et al. 2009; Xu et al. 2011; Djebali et al. 2012). An antisense transcript of *Bmal1* was also detected in a few tissues, such as subcutaneous fat, bladder, kidney, colon, lung, and potentially in duodenum, stomach, mammary gland, and ovary (Fig S6B) (Consortium 2012; Davis et al. 2018), as was also reported previously (Zhang et al. 2014). It would be of great interest to analyze whether the antisense transcript of *Bmal1* exert some functions in the mammalian circadian clock system.

Antisense transcripts of a core clock gene have also been reported in other organisms. In *Neurospora*, the sense *frequency (frq)* and antisense *(qrf)* transcripts are both located on the same chromosome and overlap almost completely (Kramer et al. 2003; Xue et al. 2014). In *Antheraea*, the sense-antisense transcripts for the homologue of *Drosophila*’s *period* (*per*) gene are located on different chromosomes (Sauman and Reppert 1996). Interestingly, human *PER2* also has an antisense transcript *(PER2AS)*, but, based on human ENCODE datasets, the transcription of *PER2AS* and *PER2* diverges from their respective TSS, most likely using a bidirectional promoter (Consortium 2012; Davis et al. 2018) (Fig. S6C). Although it is unclear whether all these sense-antisense transcripts have rhythmic and antiphasic expression patterns, the evolutionary conversation of sense-antisense pairs of a core clock gene suggests that antisense transcripts are part of a common mechanism for circadian clock regulation.

Even though the expression of *qrf* is rhythmic and anti-phasic to *frq*, similar to *Per2AS* and *Per2* (Kramer et al. 2003; Xue et al. 2014), their functions in the core clock machinery appear to be different. While the primary function of *qrf* is to regulate phase and light entrainment using both pre- and post-transcriptional regulation (Kramer et al. 2003; Xue et al. 2014), primary function of *Per2AS* appears to regulate amplitude using a pre-transcriptional mechanism (Fig. 2). Because the molecular clock circuitry is markedly more complex in mammals than *Neurospora*, it is possible that *Per2AS* acquired additional or different functions, such as its apparent ability to regulate *Bmal1 in trans*.

*Per2* is the only core clock gene whose rhythmicity, proper phase, and expression levels are all critical to sustain rhythmicity (Zheng et al. 1999; Bae et al. 2001; Chen et al. 2009). *Per2* is also under multiple layers of regulation both transcriptionally and post-transcriptionally to sustain robust rhythmicity (Yoo et al. 2005; Chen et al. 2013). This study demonstrates that *Per2AS* serves as an additional regulatory mechanism, to further ensure that *Per2’s* expression pattern stays within a certain window for sustaining robust circadian rhythmicity. It would be of great future interest to explore the functions of *Per2AS* at the tissue and organismal levels.

## Materials and Methods

### Tissue harvesting and Cell culture

Male C57BL/6J mice were maintained on a 12:12 LD cycle and fed *ad libitum*, then livers were collected at the time indicated. All the animal experiments were conducted and the protocols were approved by the Institutional Animal Care and Use Committees at the University of Texas Southwestern Medical Center.

NIH3T3, HEK293/T17, Mouse Embryonic Fibroblasts (MEFs), *PER2::LUC* MEFs, and *Bmal1-luc* cells were grown in Dulbecco’s Modified Eagle Medium (Life Tech) with 10% fetal bovine serum (FBS) (ATLANTA biologicals) at 37°C with 5% CO_2_. AML12 cells were grown in Dulbecco’s Modified Eagle Medium/F12 (1:1) with 10% FBS and 1% Insulin-Transferrin-Selenium supplement (Gibco) at 37°C with 5% CO_2_.

Chromosomal number was counted as previous reported (Nicholson et al. 2015) with minor modifications. Briefly, cell cultures were incubated in 1000 ng/ml Colcemid (Karyomax, Invitrogen) at 37°C for 3–6 hr to enrich in mitotically arrested cells. The cells were then collected by centrifugation, and pre-warmed hypotonic solution (0.075M KCl) was added dropwise to the cell pellet. After incubated for 18 min at 37°C, cells were fixed with an ice-cold 3:1 methanol:acetic acid solution for 5 min followed by centrifugation. After repeating this last step twice, fixed cells were dropped on microscope slides.

### CRISPR mutagenesis

The sgRNAs were designed by CRISPR design tool (http://crispr.mit.edu), and complementary oligonucleotides were annealed, phosphorylated and cloned into the BbsI sites of the pSpCas9(BB)-2A-GFP (Addgene plasmid # 48138) (Ran et al. 2013). After the nucleotide sequences of the plasmids were verified by Sanger sequencing, DNA was transfected into *PER2::LUC* MEFs and/or *Bmal1-luc* cells using FuGENE6 according to the manufacturer’s instructions. After 48 hrs, cells were trypsinized and sorted using FACSAria I (BD Biosciences) with GFP signals. Subsequently, cells were cultured in DMEM supplemented with 10% FBS and 1x Penicillin/Streptomycin (Gibco). When each clone reached confluency, cells were lysed with 90 μL of 50 mM NaOH and 10ul of 100 mM Tris–HCl (pH 8.0), followed by incubation at 95C for 10 min. The mutated genomic region was first amplified by PCR, then cloned into pGEM-T vector (Promega), to reveal the nucleotide sequence of each clone by Sanger sequence. All the primer sequences used in this study can be found in Supplemental Table S1.

### Real-time bioluminescence recordings and luciferase assay

Cells were first plated into 35 mm dishes and then allowed to become confluent. The medium was then replaced with DMEM supplemented with 50% horse serum (gene expression analyses) or dexamethasone (1 μM) was added to the media (real-time bioluminescence recordings) for 2 hr. After synchronizing cells, medium was changed to phenol red-free DMEM (Cellgro 90-013-PB) supplemented with 100 μM luciferrin, 10 mM HEPES pH 7.2, 1 mM sodium pyruvate, 0.035% sodium bicarbonate, 2% FBS, 1x Penicillin/Streptomycin, and 2 mM L-glutamine. Real-time bioluminescence recordings were performed using a LumiCycle (Actimetrics, Inc. Wilmette, IL). Quantification analyses was performed by JMP software (Oklejewicz et al. 2008). The recordings from the first 24 hrs were eliminated for quantitative analyses because these data are unreliable.

Luciferase assays were performed as previously reported (Kojima et al. 2010). Briefly, a mixture of plasmid DNAs containing 100 ng *Per2E2-luc* or *Bmal1-luc* firefly luciferase reporter genes, 10 ng *Renilla* luciferase reporter genes, and increasing amounts of *Per2AS* (variant2) - expressing plasmids (50, 100, 200, and 400 ng) were co-transfected in NIH3T3 cells. Luciferase activities were measured approximately 48 hrs after transfection. *Per2E2-luc* reporter gene was constructed by inserting the mouse *Per2* promoter (−83 to +156 in reference to the *Per2* TSS) into pGL4.12[*luc2CP*] (Promega), whereas *Bmal1-luc* was constructed by inserting the mouse *Bmal1* promoter (−779 to +127 in reference to the *Bmal1* TSS) to pGL4.11[*luc2P*] (Promega).

### RT-qPCR

Total RNA was extracted with TRIZOL reagent (Life Tech) according to the manufacturer’s instructions and treated with TURBO DNaseI (Life Tech). RNAs were then subjected to reverse transcription using SuperScript II (Life Tech) or High Capacity cDNA Reverse Transcription Kits (Applied Biosystems). For *Per2AS* transcripts, cDNA was synthesized using strand-specific primers (Table S1), except for Fig. 1A, in which the oligo(dT) was used. qPCR was performed using QuantiStudio 6 (Life Tech) with SYBR Power Green (Applied Biosystems). All the primer sequences used in this study can be found in Table S1.

### Rapid amplification of cDNA ends (RACE) assays

RACE assay was performed with SMARTer RACE 5’/3’ kit (Clontech) according to the manufacturer’s instructions. Total RNAs from mouse liver (C57BL/6J) harvested at ZT 4 was subjected to 5’- and 3’-RACE cDNA synthesis. Each cDNA was then amplified by *Per2AS* specific primers (Table S1) together with the Universal Primer (Clontech). The first PCR products were subsequently amplified by the nested *Per2AS* specific primers with Universal Primer (Clontech). The nested PCR products were cloned into pGEM-T vector (Promega) and the nucleotide sequence was determined by Sanger sequencing. Two independent 5’RACE products had the identical sequence, while four 3’RACE products yielded three different sequences, giving rise to the three variants of *Per2AS* (Fig. 1B, Fig. S1C).

### Northern Blotting

Total RNA (~500ug) extracted from mouse liver harvested at ZT 4, at which *Per2AS* expression is the highest (Koike et al. 2012), was subjected to poly(A)+tract isolation kit (Promega). Poly(A)^+^ enriched RNA, as well as RNA ladder (Life Tech), was further separated on a 1% agarose gel, transferred to Hybond-N^+^ membrane (Amersham), and UV cross-linked (Stratagene) using a standard protocol. The membrane was further hybridized at 68°C for overnight using PerfectHyb PLUS (Sigma) with a P^32^UTP-labeled probe that has a complementary sequence against *Per2AS* (1-1812 nt, Fig. 1B, Fig. S1C), where all the variants share its nucleotide sequences. After extensive washing of the membrane, the radioactive signal was analyzed with a Storm image analyzer (GE Healthcare).

### Subcellular fractionation

Livers from mice (C57BL/6J) were immediately rinsed with ice-cold 0.9% NaCl solution, then homogenized on ice with 5 volumes of homogenization buffer (10 mM Tris-HCl buffer, pH 7.7, containing 10 mM NaCl, 0.1 mM EGTA, 0.5 mM EDTA, 0.5 mM spermidine, 0.15 mM spermine, 0.5% Tergitol NP-10, 1 mM dithiothreitol, and 1 mM phenylmethylsulfonyl fluoride) in a Dounce homogenizer. The homogenates were filtered through two layers of cheesecloth and, following dilution with 8 tissue volumes of 2.2 M sucrose in the homogenization buffer, applied to a 10-ml cushion of 2 M sucrose in homogenization buffer and spun at 24,000 rpm for 60 min at 2 °C in a prechilled SW28 rotor. The resultant nuclear pellet as well as supernatant were subjected to RNA extraction. For NIH3T3 and MEFs, cells were lysed in an ice-cold Hypotonic Lysis buffer (10mM Tris-HCl, pH 7.4, 10mM NaCl, 3mM MgCl_2_, and 0.3% NP-40) supplemented with protease inhibitor cocktail (Sigma). After incubated for 5 min on ice with occasional pipetting, lysates were separated to nuclear and cytoplasmic fractions by centrifuging 600xg for 5 min at 4°C. RNAs were extracted from each fraction using TRIZOL reagent (Life Tech) from each fraction and subjected to RT-qPCR.

### *Per2AS* overexpression and knockdown

*Per2AS* variant 2 overexpression plasmid was generated from 5’RACE and 3’RACE products that were cloned in to pBluescript (Agilent), as well as *Per2AS* qPCR product cloned into pGEM-T (Promega). After combining 5’RACE and qPCR products in pBlueScript using Hind III and Xho I sites, the 3’RACE product was also inserted using Xho I and Kpn I sites. The full length was then transferred to pcDNA3.1-mycHis (Invitrogen) and its sequence was verified by Sanger sequencing. DNA transfection was performed with either FuGENE6 Reagent (Promega) or Lipofectamine 2000 (Life Tech) using Opti-MEM (Gibco), while gapmers were introduced via gymnosis (Stein et al. 2010) with the final concentration of 100nM (Table S1).

### Lentivirus generation and transduction

For virus production, either control or *Per2* shRNA as well as viral packaging vectors were transfected into HEK293/T17 cells according to the manufacturer’s instruction (ViraPower Lentiviral Expression System, Life Tech). Cell culture media was collected after 48 hours, ultracentrifuged at 70,000xg for 2 hours at room temperature, then resuspended in target cell-specific media. Viral media and 20μg/ml polybrene (Millipore) were added to target cells, and cells were harvested 48-72 hours after viral transduction.

### Mathematical Model

Our model of *Per2-Per2AS* interactions (Battogtokh et al. 2018) was based on an earlier model (Relogio et al. 2011) of the mammalian circadian clock, which did not take *Per2AS* into account. Our model of the basic negative feedback loop, whereby PER2:CRY inhibits BMAL:CLOCK (without additional feedback loops through REV-ERB and ROR), consisted of 15 ordinary differential equations (ODEs) with 44 parameter values (rate constants, binding constants, etc.). To illustrate some of the results of this detailed model, we present here a simpler model of the negative feedback loop (Fig. S5A) is based on Goodwin’s original model (Goodwin 1965), with an important modification suggested by Bliss, Painter and Marr (Bliss et al. 1982). *Per2AS* RNA was added to the basic Goodwin model by assuming that *Per2AS* and *Per2* mutually inhibit each other pre-transcriptionally. The ODEs describing the model are presented in Fig. S5B. The model <u>with</u> *Per2AS* consists of all four ODEs, as written. The model <u>without</u> *Per2AS* consists of the first three ODEs only, and the blue factor in the first ODE is set = 1. These two models were simulated with the XPP-AUT program (http://www.math.pitt.edu/~bard/xpp/xpp.html) using the parameter values given in Table S2. The parameter values were chosen to give oscillations with a period close to 24 hours and reasonable amplitudes and phase relations of the variables (Fig. S5C. In particular, the model with *Per2AS* was constrained to fit experimental observations in mouse liver that the level of *Per2AS* is approximately 5% of *Per2*, and that *Per2AS* and *Per2* are expressed anti-phasically (Koike et al. 2012). In Fig. S5D we compare the robustness of oscillations in the models with and without Per2AS, in terms of the range of *a*_1_ values (the maximum rate of synthesis of *Per2* mRNA) over which the models exhibit oscillations. By this measure, the model with *Per2AS* is considerably more robust than the model without *Per2AS*. In Fig. S5E, we show the domains of oscillation for both models on a two-parameter diagram: *a*_1_ and *b*_3_, where *b*_3_ is the maximum rate of degradation of phosphorylated PER2 in the nucleus. This diagram also shows a larger region of oscillations in the model with *Per2AS*. Even if we restrict our attention to the regions of “circadian” oscillations (22 < period (h) < 26) in parameter space, the model with *Per2AS* (blue lines in Fig. S5E) is noticeably more robust than the model without *Per2AS* (black lines).

## Acknowledgement

We thank Dr. Ueli Schibler (Université for Genève) for providing *Bmal1-luc* cell line, Dr. Seung-Hee Yoo (University of Texas Health Science Center at Houston) for *PER2::LUC* MEFs, and Dr. Andrew Liu (University of Florida) for *Per2* shRNA vectors (Ramanathan et al. 2014). We also thank Melissa Makris (Flow Cytometry Resource Laboratory, Department of Biomedical Sciences and Pathobiology, Center for Molecular Medicine and Infectious Diseases, Virginia-Maryland Regional College of Veterinary Medicine, Virginia Tech), Drs. Nicolaas Baudoin and Daniela Cimini (Department of Biological Sciences, Virginia Tech), and Tsubasa Toda and Akari Ueta (Department of Environmental and Life Sciences, Toyohashi University of Technology) for technical assistance. The authors also thank all the past and current members of the Green, Takahashi, Kojima laboratories for invaluable discussion. This work was supported by the Luther and Alice Hamlett Undergraduate Research Support (to C.T.S), Howard Hughes Medical Institute (J.S.T), and National Institutes of Health GM127122 (to C.B.G), and GM126223 (to S.K.). J.S.T. is an Investigator in the Howard Hughes Medical Institute.

## References

Anderson KM, Anderson DM, McAnally JR, Shelton JM, Bassel-Duby R, Olson EN. 2016. Transcription of the non-coding RNA upperhand controls Hand2 expression and heart development. Nature 539: 433–436.

Atger F, Gobet C, Marquis J, Martin E, Wang J, Weger B, Lefebvre G, Descombes P, Naef F, Gachon F. 2015. Circadian and feeding rhythms differentially affect rhythmic mRNA transcription and translation in mouse liver. Proc Natl Acad Sci USA 112: E6579–6588.

Bae K, Jin X, Maywood ES, Hastings MH, Reppert SM, Weaver DR. 2001. Differential functions of mPer1, mPer2, and mPer3 in the SCN circadian clock. Neuron 30: 525–536.

Balsalobre A, Damiola F, Schibler U. 1998. A serum shock induces circadian gene expression in mammalian tissue culture cells. Cell 93: 929–937.

Battogtokh D, Kojima S, Tyson JJ. 2018. Modeling the interactions of sense and antisense Period transcripts in the mammalian circadian clock network. PLoS computational biology 14: e1005957.

Bertone P, Stolc V, Royce TE, Rozowsky JS, Urban AE, Zhu X, Rinn JL, Tongprasit W, Samanta M, Weissman S et al. 2004. Global identification of human transcribed sequences with genome tiling arrays. Science 306: 2242–2246.

Bliss RD, Painter PR, Marr AG. 1982. Role of feedback inhibition in stabilizing the classical operon. J Theor Biol 97: 177–193.

Bond AM, Vangompel MJ, Sametsky EA, Clark MF, Savage JC, Disterhoft JF, Kohtz JD. 2009. Balanced gene regulation by an embryonic brain ncRNA is critical for adult hippocampal GABA circuitry. Nature neuroscience 12: 1020–1027.

Carninci P Kasukawa T Katayama S Gough J Frith MC Maeda N Oyama R Ravasi T Lenhard B Wells C et al. 2005. The transcriptional landscape of the mammalian genome. Science 309: 1559–1563.

Carthew RW, Sontheimer EJ. 2009. Origins and Mechanisms of miRNAs and siRNAs. Cell 136: 642–655.

Chen R, D’Alessandro M, Lee C. 2013. miRNAs are required for generating a time delay critical for the circadian oscillator. Curr Biol 23: 1959–1968.

Chen R, Schirmer A, Lee Y, Lee H, Kumar V, Yoo SH, Takahashi JS, Lee C. 2009. Rhythmic PER abundance defines a critical nodal point for negative feedback within the circadian clock mechanism. Mol Cell 36: 417–430.

Clemson CM, Hutchinson JN, Sara SA, Ensminger AW, Fox AH, Chess A, Lawrence JB. 2009. An architectural role for a nuclear noncoding RNA: NEAT1 RNA is essential for the structure of paraspeckles. Mol Cell 33: 717–726.

Consortium EP. 2012. An integrated encyclopedia of DNA elements in the human genome. Nature 489: 57–74.

Davis CA, Hitz BC, Sloan CA, Chan ET, Davidson JM, Gabdank I, Hilton JA, Jain K, Baymuradov UK, Narayanan AK et al. 2018. The Encyclopedia of DNA elements (ENCODE): data portal update. Nucleic Acids Res 46: D794–D801.

Derrien T, Johnson R, Bussotti G, Tanzer A, Djebali S, Tilgner H, Guernec G, Martin D, Merkel A, Knowles DG et al. 2012. The GENCODE v7 catalog of human long noncoding RNAs: analysis of their gene structure, evolution, and expression. Genome research 22: 1775–1789.

Dinger ME, Amaral PP, Mercer TR, Pang KC, Bruce SJ, Gardiner BB, Askarian-Amiri ME, Ru K, Solda G, Simons C et al. 2008. Long noncoding RNAs in mouse embryonic stem cell pluripotency and differentiation. Genome research 18: 1433–1445.

Djebali S, Davis CA, Merkel A, Dobin A, Lassmann T, Mortazavi A, Tanzer A, Lagarde J, Lin W, Schlesinger F et al. 2012. Landscape of transcription in human cells. Nature 489: 101–108.

Engreitz JM, Haines JE, Perez EM, Munson G, Chen J, Kane M, McDonel PE, Guttman M, Lander ES. 2016. Local regulation of gene expression by lncRNA promoters, transcription and splicing. Nature 539: 452–455.

Faghihi MA, Wahlestedt C. 2009. Regulatory roles of natural antisense transcripts. Nat Rev Mol Cell Biol 10: 637–643.

Fang B, Everett LJ, Jager J, Briggs E, Armour SM, Feng D, Roy A, Gerhart-Hines Z, Sun Z, Lazar MA. 2014. Circadian enhancers coordinate multiple phases of rhythmic gene transcription in vivo. Cell 159: 1140–1152.

Feng J, Bi C, Clark BS, Mady R, Shah P, Kohtz JD. 2006. The Evf-2 noncoding RNA is transcribed from the Dlx-5/6 ultraconserved region and functions as a Dlx-2 transcriptional coactivator. Genes Dev 20: 1470–1484.

Goodman AJ, Daugharthy ER, Kim J. 2013. Pervasive antisense transcription is evolutionarily conserved in budding yeast. Mol Biol Evol 30: 409–421.

Goodwin BC. 1965. Oscillatory behavior in enzymatic control processes. Adv Enzyme Regul 3: 425–438.

Groff AF, Sanchez-Gomez DB, Soruco MML, Gerhardinger C, Barutcu AR, Li E, Elcavage L, Plana O, Sanchez LV, Lee JC et al. 2016. In Vivo Characterization of Linc-p21 Reveals Functional cis-Regulatory DNA Elements. Cell reports 16: 2178–2186.

Guttman M, Amit I, Garber M, French C, Lin MF, Feldser D, Huarte M, Zuk O, Carey BW, Cassady JP et al. 2009. Chromatin signature reveals over a thousand highly conserved large non-coding RNAs in mammals. Nature 458: 223–227.

Halley P, Kadakkuzha BM, Faghihi MA, Magistri M, Zeier Z, Khorkova O, Coito C, Hsiao J, Lawrence M, Wahlestedt C. 2014. Regulation of the apolipoprotein gene cluster by a long noncoding RNA. Cell reports 6: 222–230.

He B, Nohara K, Park N, Park YS, Guillory B, Zhao Z, Garcia JM, Koike N, Lee CC, Takahashi JS et al. 2016. The Small Molecule Nobiletin Targets the Molecular Oscillator to Enhance Circadian Rhythms and Protect against Metabolic Syndrome. Cell metabolism 23: 610–621.

He Y, Vogelstein B, Velculescu VE, Papadopoulos N, Kinzler KW. 2008. The antisense transcriptomes of human cells. Science 322: 1855–1857.

Janich P, Arpat AB, Castelo-Szekely V, Lopes M, Gatfield D. 2015. Ribosome profiling reveals the rhythmic liver translatome and circadian clock regulation by upstream open reading frames. Genome research 25: 1848–1859.

Johnsson P, Lipovich L, Grander D, Morris KV. 2014. Evolutionary conservation of long non-coding RNAs; sequence, structure, function. Biochim Biophys Acta 1840: 1063–1071.

Katayama S, Tomaru Y, Kasukawa T, Waki K, Nakanishi M, Nakamura M, Nishida H, Yap CC, Suzuki M, Kawai J et al. 2005. Antisense transcription in the mammalian transcriptome. Science 309: 1564–1566.

Khaitovich P, Kelso J, Franz H, Visagie J, Giger T, Joerchel S, Petzold E, Green RE, Lachmann M, Paabo S. 2006. Functionality of intergenic transcription: an evolutionary comparison. PLoS Genet 2: e171.

Khorkova O, Myers AJ, Hsiao J, Wahlestedt C. 2014. Natural antisense transcripts. Human molecular genetics 23: R54–63.

Koike N, Yoo SH, Huang HC, Kumar V, Lee C, Kim TK, Takahashi JS. 2012. Transcriptional Architecture and Chromatin Landscape of the Core Circadian Clock in Mammals. Science 338: 349.

Kojima S, Gatfield D, Esau CC, Green CB. 2010. MicroRNA-122 modulates the rhythmic expression profile of the circadian deadenylase Nocturnin in mouse liver. PLoS One 5: e11264.

Kramer C, Loros JJ, Dunlap JC, Crosthwaite SK. 2003. Role for antisense RNA in regulating circadian clock function in Neurospora crassa. Nature 421: 948–952.

Lee JE, Bennett CF, Cooper TA. 2012. RNase H-mediated degradation of toxic RNA in myotonic dystrophy type 1. Proc Natl Acad Sci USA 109: 4221–4226.

Lee JS, Mendell JT. 2020. Antisense-Mediated Transcript Knockdown Triggers Premature Transcription Termination. Mol Cell 77: 1044–1054 e1043.

Lee JT, Davidow LS, Warshawsky D. 1999. Tsix, a gene antisense to Xist at the X-inactivation centre. Nat Genet 21: 400–404.

Littleton ES, Kojima S. 2020. Genome-wide correlation analysis reveals <em>Rorc</em> as potential amplitude regulator of circadian transcriptome output. bioRxiv: 2020.2006.2019.161307.

Lowrey PL, Takahashi JS. 2004. Mammalian circadian biology: elucidating genome-wide levels of temporal organization. Annual review of genomics and human genetics 5: 407–441.

Menet JS, Rodriguez J, Abruzzi KC, Rosbash M. 2012. Nascent-Seq reveals novel features of mouse circadian transcriptional regulation. eLife 1: e00011.

Mercer TR, Dinger ME, Mattick JS. 2009. Long non-coding RNAs: insights into functions. Nat Rev Genet 10: 155–159.

Modarresi F, Faghihi MA, Lopez-Toledano MA, Fatemi RP, Magistri M, Brothers SP, van der Brug MP, Wahlestedt C. 2012. Inhibition of natural antisense transcripts in vivo results in gene-specific transcriptional upregulation. Nat Biotechnol 30: 453–459.

Morf J, Rey G, Schneider K, Stratmann M, Fujita J, Naef F, Schibler U. 2012. Cold-inducible RNA-binding protein modulates circadian gene expression posttranscriptionally. Science 338: 379–383.

Morris KV, Santoso S, Turner AM, Pastori C, Hawkins PG. 2008. Bidirectional transcription directs both transcriptional gene activation and suppression in human cells. PLoS Genet 4: e1000258.

Nicholson JM, Macedo JC, Mattingly AJ, Wangsa D, Camps J, Lima V, Gomes AM, Doria S, Ried T, Logarinho E et al. 2015. Chromosome mis-segregation and cytokinesis failure in trisomic human cells. eLife 4.

Oklejewicz M, Destici E, Tamanini F, Hut RA, Janssens R, van der Horst GT. 2008. Phase resetting of the mammalian circadian clock by DNA damage. Curr Biol 18: 286–291.

Panda AC, Grammatikakis I, Munk R, Gorospe M, Abdelmohsen K. 2017. Emerging roles and context of circular RNAs. Wiley interdisciplinary reviews RNA 8.

Pang KC, Frith MC, Mattick JS. 2006. Rapid evolution of noncoding RNAs: lack of conservation does not mean lack of function. Trends Genet 22: 1–5.

Phatnani HP, Greenleaf AL. 2006. Phosphorylation and functions of the RNA polymerase II CTD. Genes Dev 20: 2922–2936.

Ponjavic J, Ponting CP, Lunter G. 2007. Functionality or transcriptional noise? Evidence for selection within long noncoding RNAs. Genome research 17: 556–565.

Ramanathan C, Xu H, Khan SK, Shen Y, Gitis PJ, Welsh DK, Hogenesch JB, Liu AC. 2014. Cell type-specific functions of period genes revealed by novel adipocyte and hepatocyte circadian clock models. PLoS Genet 10: e1004244.

Ran FA, Hsu PD, Wright J, Agarwala V, Scott DA, Zhang F. 2013. Genome engineering using the CRISPR-Cas9 system. Nature protocols 8: 2281–2308.

Rhind N, Chen Z, Yassour M, Thompson DA, Haas BJ, Habib N, Wapinski I, Roy S, Lin MF, Heiman DI et al. 2011. Comparative functional genomics of the fission yeasts. Science 332: 930–936.

Rosok O, Sioud M. 2004. Systematic identification of sense-antisense transcripts in mammalian cells. Nat Biotechnol 22: 104–108.

Sato TK, Panda S, Miraglia LJ, Reyes TM, Rudic RD, McNamara P, Naik KA, FitzGerald GA, Kay SA, Hogenesch JB. 2004. A functional genomics strategy reveals Rora as a component of the mammalian circadian clock. Neuron 43: 527–537.

Sauman I, Reppert SM. 1996. Circadian clock neurons in the silkmoth Antheraea pernyi: novel mechanisms of Period protein regulation. Neuron 17: 889–900.

Schmutz I, Ripperger JA, Baeriswyl-Aebischer S, Albrecht U. 2010. The mammalian clock component PERIOD2 coordinates circadian output by interaction with nuclear receptors. Genes Dev 24: 345–357.

Shearman LP, Sriram S, Weaver DR, Maywood ES, Chaves I, Zheng B, Kume K, Lee CC, van der Horst GT, Hastings MH et al. 2000. Interacting molecular loops in the mammalian circadian clock. Science 288: 1013–1019.

Sleutels F, Zwart R, Barlow DP. 2002. The non-coding Air RNA is required for silencing autosomal imprinted genes. Nature 415: 810–813.

Smilinich NJ, Day CD, Fitzpatrick GV, Caldwell GM, Lossie AC, Cooper PR, Smallwood AC, Joyce JA, Schofield PN, Reik W et al. 1999. A maternally methylated CpG island in KvLQT1 is associated with an antisense paternal transcript and loss of imprinting in Beckwith-Wiedemann syndrome. Proc Natl Acad Sci USA 96: 8064–8069.

Sopher BL, Ladd PD, Pineda VV, Libby RT, Sunkin SM, Hurley JB, Thienes CP, Gaasterland T, Filippova GN, La Spada AR. 2011. CTCF regulates ataxin-7 expression through promotion of a convergently transcribed, antisense noncoding RNA. Neuron 70: 1071–1084.

Stein CA, Hansen JB, Lai J, Wu S, Voskresenskiy A, Hog A, Worm J, Hedtjarn M, Souleimanian N, Miller P et al. 2010. Efficient gene silencing by delivery of locked nucleic acid antisense oligonucleotides, unassisted by transfection reagents. Nucleic Acids Res 38: e3.

Sun M, Hurst LD, Carmichael GG, Chen J. 2006. Evidence for variation in abundance of antisense transcripts between multicellular animals but no relationship between antisense transcriptionand organismic complexity. Genome research 16: 922–933.

Takahashi JS. 2017. Transcriptional architecture of the mammalian circadian clock. Nat Rev Genet 18: 164–179.

Takahashi JS, Hong HK, Ko CH, McDearmon EL. 2008. The genetics of mammalian circadian order and disorder: implications for physiology and disease. Nat Rev Genet 9: 764–775.

Villegas VE, Rahman MF, Fernandez-Barrena MG, Diao Y, Liapi E, Sonkoly E, Stahle M, Pivarcsi A, Annaratone L, Sapino A et al. 2014. Identification of novel non-coding RNA-based negative feedback regulating the expression of the oncogenic transcription factor GLI1. Mol Oncol 8: 912–926.

Vollmers C, Schmitz RJ, Nathanson J, Yeo G, Ecker JR, Panda S. 2012. Circadian oscillations of protein-coding and regulatory RNAs in a highly dynamic mammalian liver epigenome. Cell metabolism 16: 833–845.

Wanowska E, Kubiak MR, Rosikiewicz W, Makalowska I, Szczesniak MW. 2018. Natural antisense transcripts in diseases: From modes of action to targeted therapies. Wiley interdisciplinary reviews RNA 9.

Wight M, Werner A. 2013. The functions of natural antisense transcripts. Essays Biochem 54: 91–101.

Xu Z, Wei W, Gagneur J, Clauder-Munster S, Smolik M, Huber W, Steinmetz LM. 2011. Antisense expression increases gene expression variability and locus interdependency. Molecular systems biology 7: 468.

Xu Z, Wei W, Gagneur J, Perocchi F, Clauder-Munster S, Camblong J, Guffanti E, Stutz F, Huber W, Steinmetz LM. 2009. Bidirectional promoters generate pervasive transcription in yeast. Nature 457: 1033–1037.

Xue Z, Ye Q, Anson SR, Yang J, Xiao G, Kowbel D, Glass NL, Crosthwaite SK, Liu Y. 2014. Transcriptional interference by antisense RNA is required for circadian clock function. Nature 514: 650–653.

Yassour M, Pfiffner J, Levin JZ, Adiconis X, Gnirke A, Nusbaum C, Thompson DA, Friedman N, Regev A. 2010. Strand-specific RNA sequencing reveals extensive regulated long antisense transcripts that are conserved across yeast species. Genome Biol 11: R87.

Yoo SH, Ko CH, Lowrey PL, Buhr ED, Song EJ, Chang S, Yoo OJ, Yamazaki S, Lee C, Takahashi JS. 2005. A noncanonical E-box enhancer drives mouse Period2 circadian oscillations in vivo. Proc Natl Acad Sci U S A 102: 2608–2613.

Yoo SH, Yamazaki S, Lowrey PL, Shimomura K, Ko CH, Buhr ED, Siepka SM, Hong HK, Oh WJ, Yoo OJ et al. 2004. PERIOD2::LUCIFERASE real-time reporting of circadian dynamics reveals persistent circadian oscillations in mouse peripheral tissues. Proc Natl Acad Sci USA 101: 5339–5346.

Zhang R, Lahens NF, Ballance HI, Hughes ME, Hogenesch JB. 2014. A circadian gene expression atlas in mammals: implications for biology and medicine. Proc Natl Acad Sci U S A 111:16219–16224.

Zhao J, Sun BK, Erwin JA, Song JJ, Lee JT. 2008. Polycomb proteins targeted by a short repeat RNA to the mouse X chromosome. Science 322: 750–756.

Zheng B, Larkin DW, Albrecht U, Sun ZS, Sage M, Eichele G, Lee CC, Bradley A. 1999. The mPer2 gene encodes a functional component of the mammalian circadian clock. Nature 400: 169–173.

