## Supplementary material for "Natural Antisense Transcript of *Period2, Per2AS*, regulates the amplitude of the mouse circadian clock": Suppl Figures

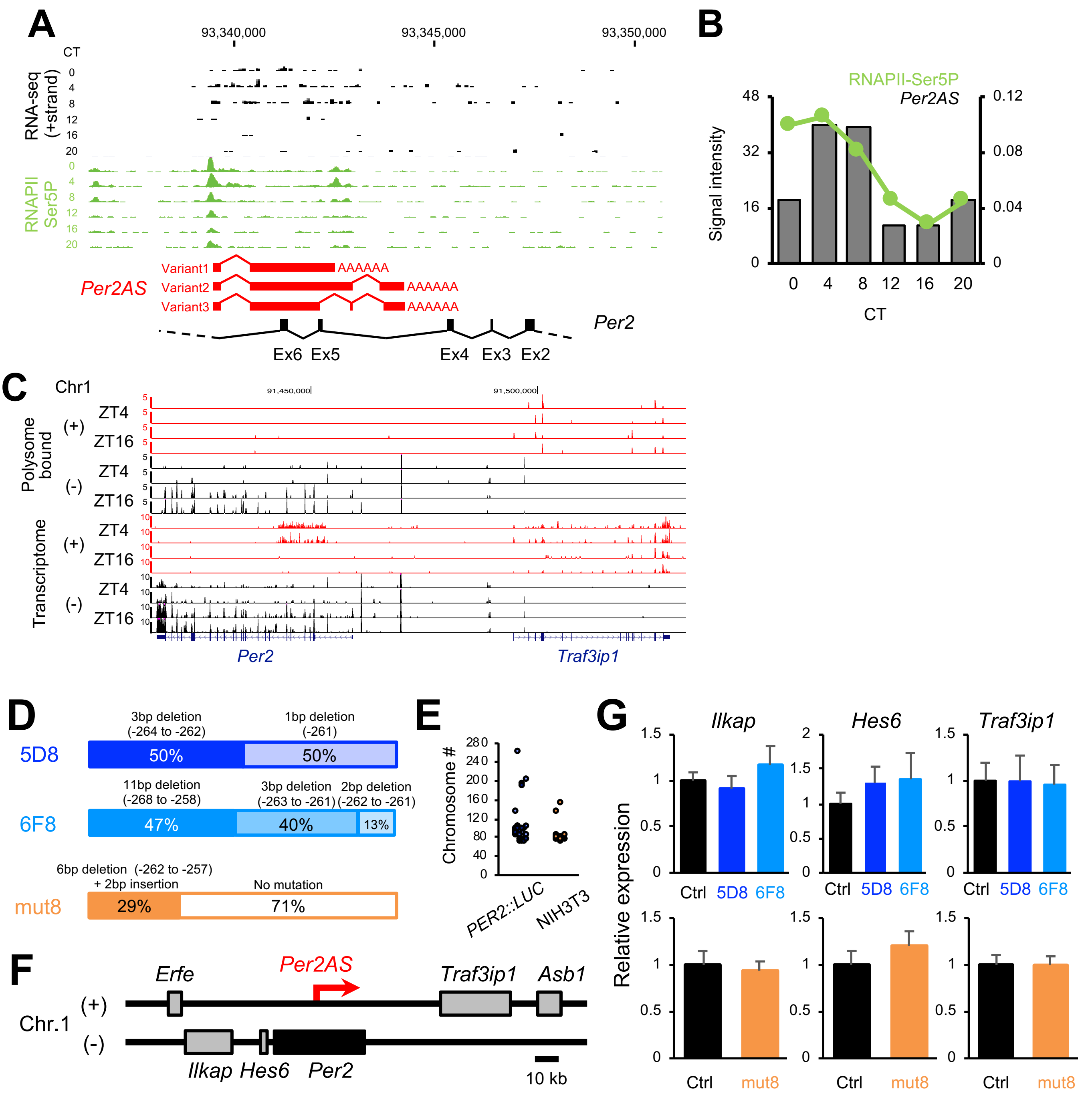


**Supplementary Figure 1: Characteristics of *Per2AS* and its mutant clones.** A) Genome browser view of the patterns of *Per2AS* expression as well as RNAPII-Ser5P accumulation on chromosome 1 (mm9) (Koike et al., 2012). Genomic structures of *Per2* and *Per2AS* are also depicted. B) Quantifications of the level of RNAPII-Ser5P accumulation (green) at the *Per2AS* TSS. Red bar graph represents *Per2AS* expression patterns (Koike et al., 2012). C) Genome browser view on chromosome 1 (mm10) for both polysome-bound transcripts (top) and transcriptome (bottom) (Janich et al., 2015). D) Genotypes and allele frequencies of *Per2AS* mutant clones. E) Distribution of chromosome numbers in NIH3T3 cells (n=28) and *PER2::LUC* MEFs (n=29). F) Genomic structure near *Per2AS*. G) The expression level of *Ilkap*, *Hes6*, and *Traf3ip1* in the *Per2AS* mutants. Data represent Mean ± SEM (n=8-13).

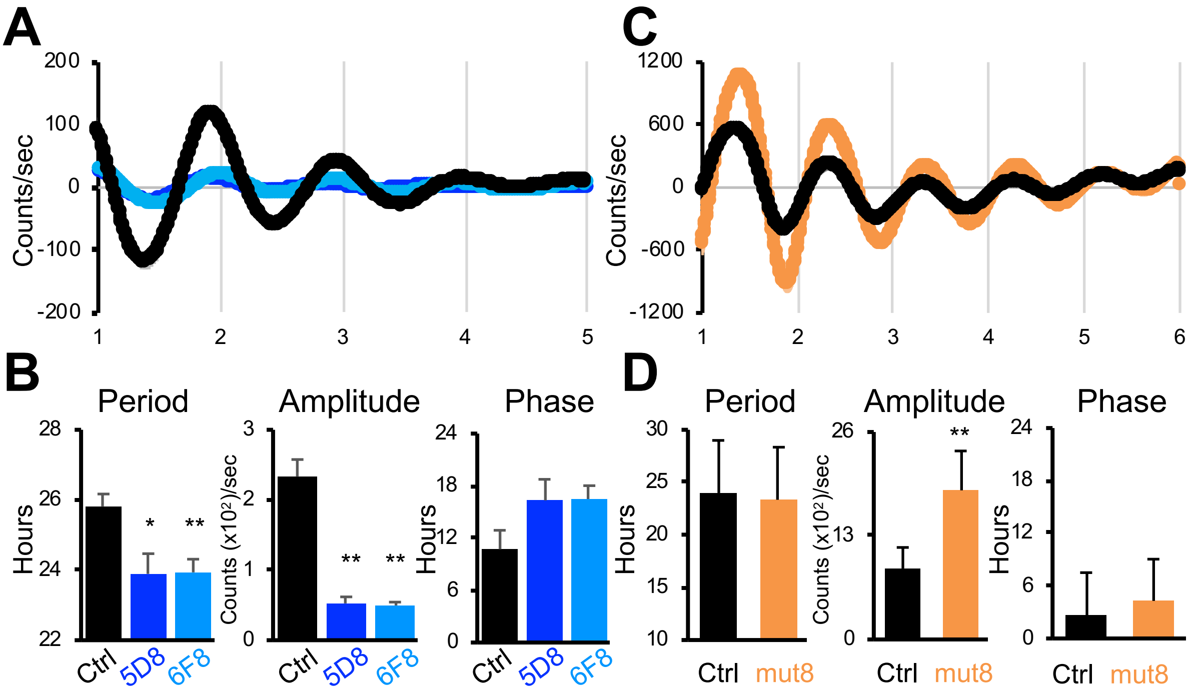


**Supplementary Figure 3. Detrended data of bioluminescent output from the *Per2AS* mutants.** A, C) *PER2::LUC* MEFs (A) or *Bmal1-luc* (C). Bold lines represent the mean, while shaded areas represent the SEM. B, D) Period (left), amplitude (middle), and phase (right) of bioluminescent output calculated from Fig. S3A or S3C. All the data represent Mean ± SEM. *; $p<0.05$, **; $p<0.005$ (Student t-test). *PER2::LUC* MEFs; WT: n=10, 5D8: n=6, 6F8: n=8, *Bmal1-luc;* WT: n=9, mut8: n=9

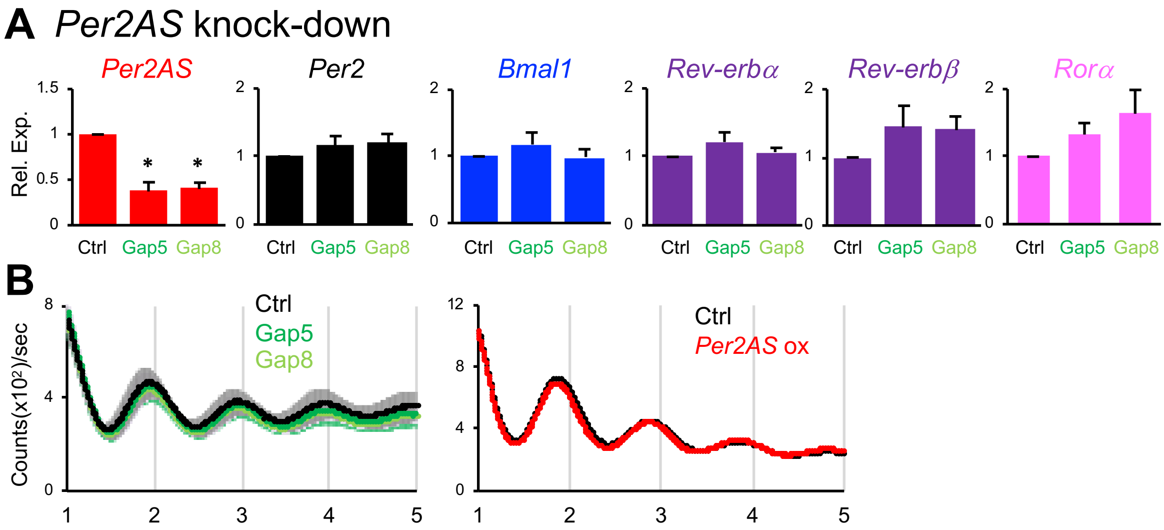


**Supplementary Figure 4. Effects of *Per2AS* transcripts in *PER2::LUC* MEFs.** A) Relative expression levels of *Per2AS*, *Per2*, and *Bmal1*, normalized by *36B4* upon *Per2AS* knock-down (n=4). All the data represent Mean $\pm$ SEM. *; $p<0.05$. B) Bioluminescent output from *Bmal1-luc* cells upon knock-down (n=10) or overexpression (n=9) of *Per2AS*. Bold lines represent the mean, while shaded areas represent the SEM.

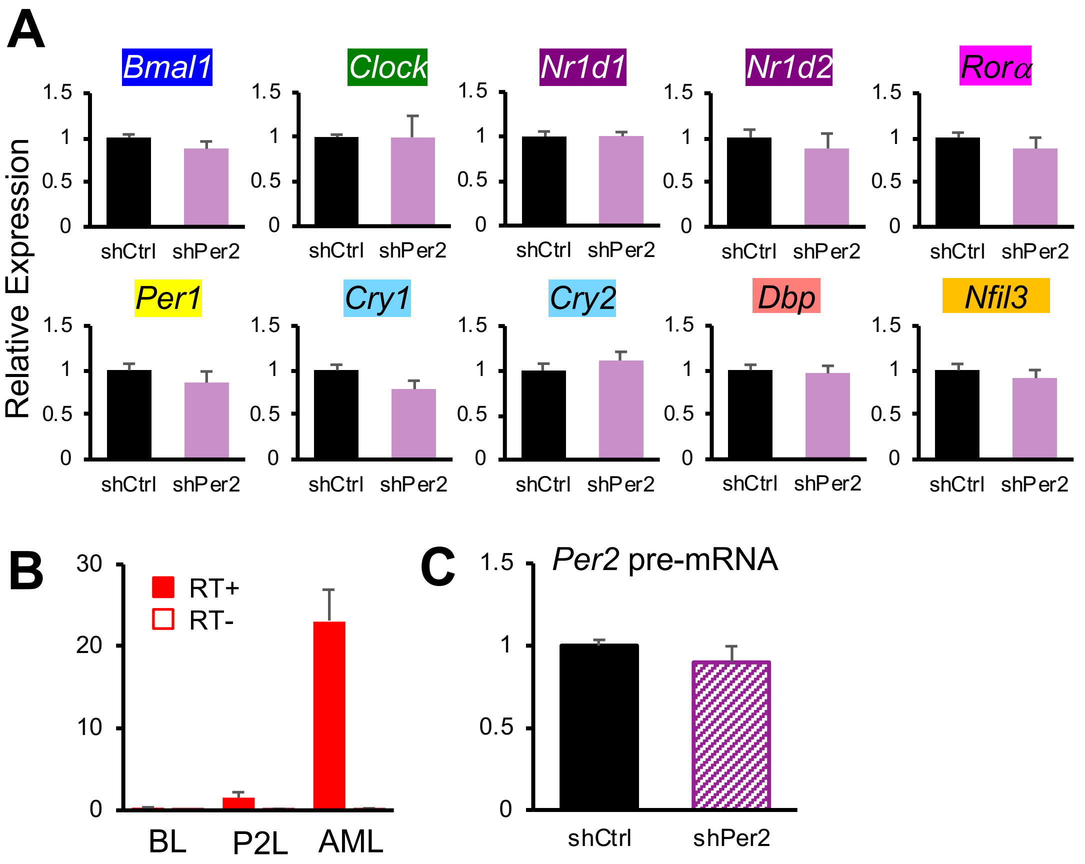


**Supplementary Figure 5. Effects of *Per2* knock-down in AML12 cell.** (A) Relative expression levels of core clock genes normalized by *36B4* (n=4-5). (B) Comparison of the level of *Per2AS* normalized by *36B4* in various cells (*Bmal1-luc* (BL) cells: n=13, PER2::LUC (P2L) MEFs: n=5, AML12 cells: n=5). (C) Relative expression level of *Per2* pre-mRNA normalized by *36B4* (n=4-5). All the data represent Mean ± SEM.

A

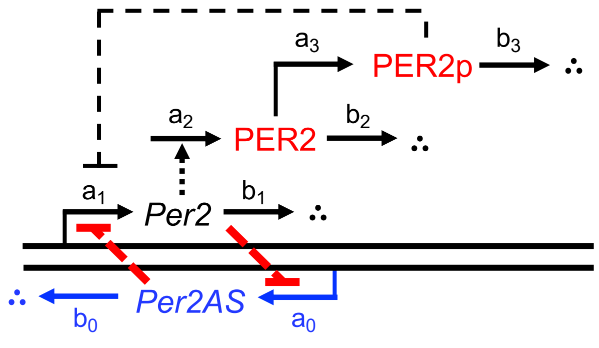


B

$$\frac{d[Per2]}{dt} = \frac{a_1}{1+[PER2p]} * \frac{K_{AS}}{K_{AS}+[Per2AS]} - b_1[Per2]$$

$$\frac{d[PER2]}{dt} = a_2[Per2] - (b_2+a_3)[PER2]$$

$$d[PER2p] = a_3[PER2] - \frac{b_3[PER2p]}{K+[PER2p]}$$

$$\frac{d[Per2AS]}{dt} = a_0 * \frac{K_S}{K_S+[Per2]} - b_0[Per2AS]$$

C

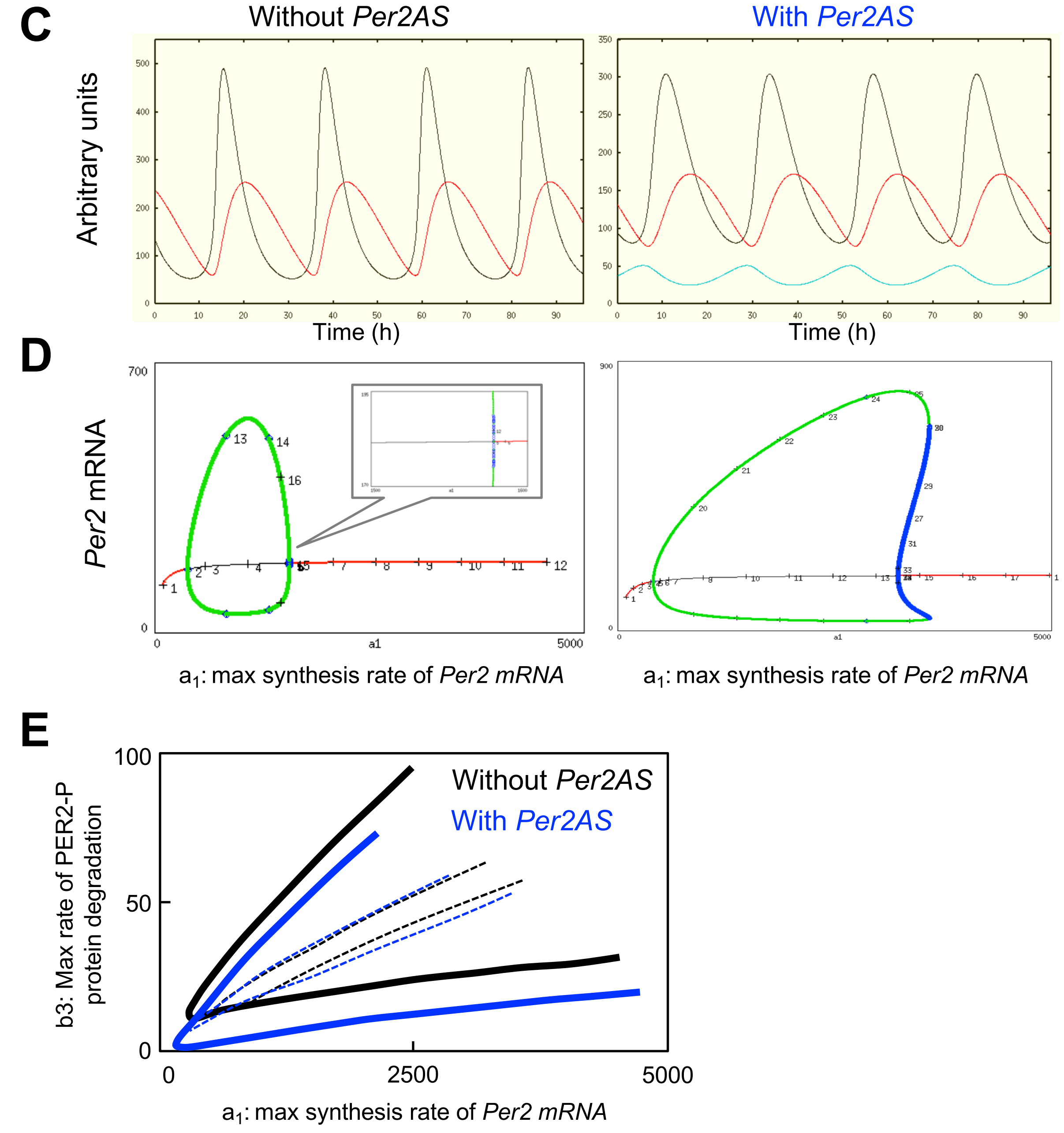

**Fig. S5: A simple mathematical formulation of the pre-transcriptional model of *Per2AS* function.** A) Wiring diagram. *Per2AS*, *Per2AS* RNA; *Per2*, *Per2* mRNA; PER2, PER2 protein; PER2p, phosphorylated PER2 protein. The red bars indicate presumed transcriptional interference between *Per2* and *Per2AS*. The $a_i$'s and $b_i$'s are rate constants for the associated reactions. B) Dynamic equations describing the model in panel A. The base model, without *Per2AS*, consists of the first three equations only, with the blue factor set = 1. C) Time-course plot of the levels of *Per2AS* (blue), *Per2* (black), and total PER2 protein (red) (i.e., PER2 + PER2p), for the parameter values listed in Table S2. In the model without *Per2AS* (left), the period is ~23 hours and the phase difference between the peaks of *Per2* and PER2 is ~4.3 hours. In the model with *Per2AS* (right), the period is ~22 hours and the phase difference between the minimum of *Per2* and the maximum of *Per2AS* is ~3 hours. D) Oscillatory region with respect to $a_1$ (the maximum rate of *Per2* mRNA synthesis) in the absence (left) and presence (right) of *Per2AS*. Without *Per2AS*, stable oscillations (green curves trace the maximum and minimum values of *Per2* mRNA) occur in the interval $391 < a_1 < 1578$, and with Per2AS, oscillations extend from $a_1 = 416$ to 3234. The Hopf bifurcations at 1578 and 3234 are subcritical (see insets), with cyclic fold bifurcations at $a_1 = 1579$ (without *Per2AS*) and at $a_1 = 3595$ (with *Per2AS*), respectively. E) Oscillatory domain in dependence on two parameters: $a_1$ and $b_3$ (the maximum rate of degradation of PER2p. The red (blue) curve shows the locus of Hopf bifurcations for the model without (with) *Per2AS*. The red and blue dashed lines indicate the loci of periodic solutions of period = 22 (uppermost) and 26 (lowermost) hours, i.e., the regions of circadian oscillations for both models. The circadian region for the model with *Per2AS* is noticeably larger than for the model without *Per2AS*.

Supplemental Fig 6. Mosig et al.,

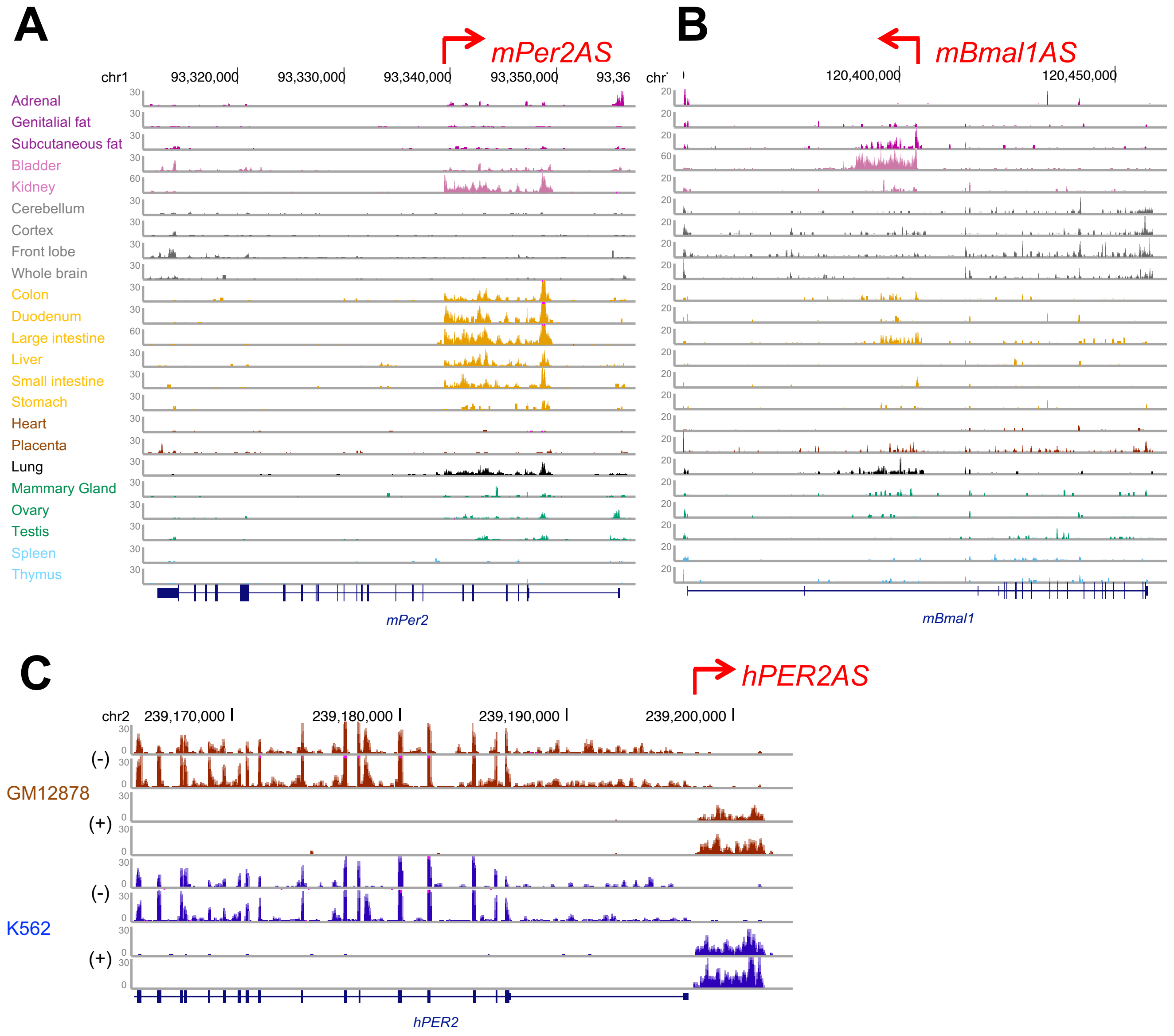


**Fig. S6: Genome browser view of antisense transcripts for m*Per2* (A), m*Bmal1* (B), and h*PER2* (C).** Data were retrieved from ENCODE project at UCSC (Consortium 2012; Davis et al. 2018). For m*Per2AS* and m*Bmal1*, only the antisense strand is displayed. The structure and location of sense genes are also displayed below the coverage plots (human: GRCh37/hg19, mouse: NCBI37/mm9).

Table S2. Description of parameters in the model (Fig. S5).

| Parameter | Description | Value | |
|---|---|---|---|
| | | Without *Per2AS* | With *Per2AS* |
| $a_1$ | Max synthesis rate of *Per2* mRNA | 780 | 600 |
| $b_1$ | Rate constant of *Per2* mRNA degradation | 0.26 | 0.2 |
| $a_2$ | Synthesis rate of PER2 protein per unit *Per2* mRNA | 0.156 | 0.12 |
| $b_2$ | Rate constant of PER2 protein degradation | 0.078 | 0.06 |
| $a_3$ | Phosphorylation rate of PER2 protein | 0.156 | 0.12 |
| $b_3$ | Max rate of PER2-P protein degradation | 19.5 | 15 |
| K | Michaelis constant for PER2-P degradation | 1 | 1 |
| $a_0$ | Max synthesis rate of *Per2AS* | NA | 200 |
| $b_0$ | Rate constant of *Per2AS* degradation | NA | 0.2 |
| $K_S$ | Dissociation (50% inhibition) constant | NA | 1 |
| $K_{AS}$ | Dissociation (50% inhibition) constant | NA | 10 |
